# A fluorescent folding reporter uncovers myosin misfolding as a driver of Hypertrophic Cardiomyopathy

**DOI:** 10.1101/2025.02.10.636595

**Authors:** Renato Arnese, Philipp Julian Dexheimer, Beatriz Escobar-Doncel, Lotta Krüger, Govinda Adhikari, Nina Krpan, Daria Sehr, Juliane Kley, Luiza Deszcz, Anton Meinhart, Janine Kirstein, Tim Clausen

## Abstract

Hypertrophic cardiomyopathy (HCM) is a fatal genetic disorder causing the thickening of ventricular walls in the heart muscle. While certain mutations in cardiac myosin deregulate ATPase activity, the pathology mechanism of most HCM mutations is not known. Here, by designing a fluorescent reporter to monitor myosin folding in cells, we uncovered a distinct class of HCM mutations that cause graded defects in myosin maturation. Using *C. elegans* as a disease model, we found that folding deficient HCM variants cause myofilament disruption, impaired motility, and reduced lifespan. Dietary restrictions resulted in a near-complete recovery from these detrimental defects by activating autophagy pathways through insulin/TOR signaling. In conclusion, our study identifies myosin misfolding as an important driver of HCM, revealing therapeutic opportunities to counteract muscle protein disorders.

## Introduction

All cells rely on a complex proteostasis network to ensure proper protein function^1^. When this network becomes overwhelmed, for example under extreme stress situations or with aging, proteins can misfold and associate into non-native, aggregation-prone entities^2,3^. Such proteotoxic stress is a hallmark of protein misfolding diseases, particularly in post-mitotic cells that cannot undergo replacement, as exemplified by neurodegenerative disorders like Alzheimer’s and Parkinson’s disease^4^.

Muscle cells are also susceptible to protein misfolding, due to mechanical stress, limited regeneration, and their specialized protein composition. In particular, myosin intricate fold and its sheer abundance - making up to 20% of total protein content in human myocytes^5^ - present a substantial challenge to the proteostasis network^6^. To ensure proper myosin folding, muscle cells express the myosin chaperone UNC-45^7,8^. Recently, defects in the myosin/UNC-45 interaction caused by a point mutation in embryonic myosin^9^ was linked to Freeman-Sheldon syndrome^10^, a rare developmental myopathy. Moreover, mutations in UNC-45 itself^11^ and the disaggregase p97^12^ have been linked to myopathies, suggesting that the intricate myocytes proteostasis network could be a muscle vulnerability.

Hypertrophic cardiomyopathy (HCM) is the most prevalent cardiac muscle disease, affecting 1 in 200 people^13^. It manifests in thickening of the ventricular walls of the heart, arrhythmia and ultimately cardiac failure, and is linked mostly to mutations in beta-cardiac myosin. Previous studies suggest that HCM mutations deregulate ATPase activity of myosin^14^ or destabilize its auto-inhibited conformation, known as super-relaxed (SRX) state^15-18^, causing contractile dysregulation in cardiomyocytes^19^. Notably, the majority of HCM disease mutations are not part of any protein motif linked to ATPase activity, actin binding or maintaining the latent SRX state of cardiac myosin. Rather, they are distributed over the entire motor domain, hinting at a broader perturbation of myosin function^20^.

To address the role of myosin proteostasis in HCM, we developed a reporter of myosin folding and screened multiple variants linked to the disease, identifying three classes of myosin mutation with graded folding defects. Modelling of these mutations in *C. elegans* unveiled myosin aggregation, motility defects and reduced lifespan, which can be exacerbated by defective chaperoning and seeds of aggregation. Strikingly, this phenotype can be reverted by caloric restriction or insulin/TOR pathway inhibition in an autophagy-dependent fashion, highlighting the role of efficient proteostasis in cardiomyopathies.

## RESULTS

### HCM mutations cause graded folding defects in myosin

To investigate the disease mechanism of HCM-linked myosin mutations we used *C. elegans* MHC-B - the major myosin isoform in body wall muscles - as our model system. The motor domain of nematode myosin shares 57% sequence identity with human cardiac myosin MYH7, and out of the 455 conserved residues, 194 are found mutated in HCM patients (**Fig. S1A**). Contrary to human MYH7, which cannot be produced in non-muscle cells, the folding of MHC-B can be recapitulated in insect cells by co-expressing its cognate chaperone UNC-45, highlighting the value of nematode myosin as myopathy model^9,21^.

To explore the putative destabilizing effect of HCM mutations on muscle myosin, we developed a fluorescent reporter monitoring the maturation of myosin within a cell. For this purpose, we engineered a split-fluorescence protein (split-FP) based on the modelled complex between the MHC-B motor domain (residues 1-819) and its essential light chain (ELC) (**Fig. 1A**). Given the proximity of the protein termini, we fused one fragment of mCherry (residues 160–239) to the C-terminus of the MHC-B motor domain, while the other portion of mCherry (residues 1–159) was attached to the C-terminus of ELC (**Fig. 1B**). When co-expressing the two proteins with UNC-45, the fluorescence signal should thus only appear when the folded MHC-B myosin interacts with the ELC, aligning the two halves of split-mCherry. Furthermore, we attached GFP to the N-terminus of MHC-B, to measure the total amount of the myosin reporter and account for variations in transcriptional and translational efficiency (**Fig. 1C**).

**Figure 1:**
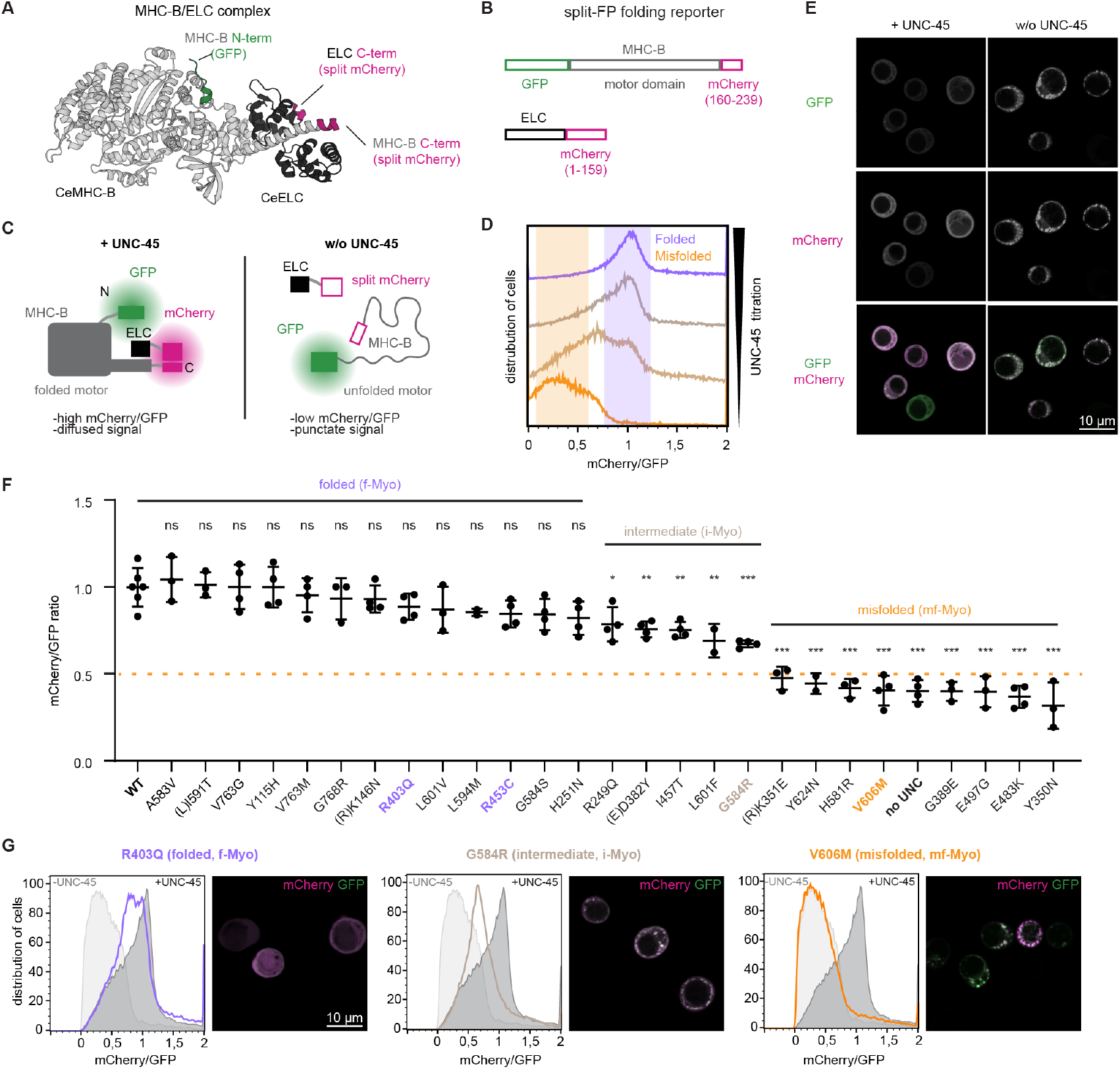
HCM mutations cause graded folding defects in myosin. **A)** Modelled MHC-B:ELC complex guiding the design of the folding reporter. **B)** Vectors of folding reporter. **C)** Schematic of the co-expression of myosin and the chaperone UNC-45 in Hi5 insect cells: with UNC-45, we expect high mCherry signal, and a diffused distribution. Without UNC-45, a low mCherry signal, and aggregated protein. **D)** Flow-cytometry analysis of Hi5 cells expressing the myosin folding reporter at increasing amounts of UNC-45 (orange, misfolded myosin; lilac, folded myosin; brown, intermediate state). **E)** Confocal images of Hi5 cells expressing the myosin folding reporter in presence and absence of UNC-45. **F)** Median of mCherry/ GFP (CG) ratio of myosin variants normalized to wildtype (WT). Amino acids in parenthesis list non-conserved residues of *C. elegans* MHC-B. Based on the CG ratio of wildtype (WT, CG 1.0) and misfolded myosin (no UNC-45, CG 0.4), HCM mutants were grouped into folded (f-Myo, CG 0.8-1.0), intermediate (i-Myo, CG 0.5-0.8) and misfolded myosins (mf-Myo, CG ≤0.5). Statistical significance against WT MHC-B is indicated after t-test. **G)** Flow-cytometry profile and aggregation behaviour of representative f-Myo (R403Q), i-Myo (G584R) and mf-Myo (V606M) variants in Hi5 cells.

To test the functionality of the reporter, we first monitored myosin folding at different UNC-45 levels. At higher levels of UNC-45 expression, we saw an increase in the mCherry signal (folded myosin) relative to GFP (total myosin), as measured by flow-cytometry (**Fig. 1D**). Moreover, we observed that the folding reporter showed a homogenous distribution in insect cells when co-expressed with UNC-45, while it aggregated in discrete foci when the chaperone was absent (**Fig. 1E**). The distribution of folded and aggregated myosin was confirmed by Western Blot analyses, measuring MHC-B levels in the soluble and insoluble fractions of insect cell lysates (**Fig. S1B**). To probe the folding reporter in a functional assay, we co-expressed MHC-B and UNC-45 mutations known to interfere with myosin folding. For MHC-B, we used mutations (F577A, H581A, Y582A) that abrogate the interaction with the UNC-45 chaperone and cause the muscle protein to aggregate in cells^9^. For UNC-45, we used a variant lacking the domain for myosin binding (ΔUCS)^22^. A strong decrease in the mCherry/GFP ratio indicated that myosin folding was indeed impaired by each of the mutations (**Fig. S1C-D**). Together, these data show that the split-FP reporter reflects the relative amounts of folded and misfolded myosin in the cell, allowing us to study the folding of its motor domain in a non-intrusive, quantitative manner.

To test the relevance of myosin misfolding as myopathy driver, we introduced disease-linked point mutations of human β-cardiac myosin MYH7 into *C. elegans* MHC-B and generated 26 folding reporter constructs (**Table S1**, numbering of residues according to human MYH7). Myosin variants included well-characterized HCM mutations that interfere with ATPase function or SRX formation, mutations found at the myosin/UNC-45 interface, and disease mutations with unknown mechanism. Flow-cytometry analysis revealed that HCM mutations caused a continuum of folding defects, ranging from little or no impact on the folded state to mutant proteins that were completely misfolded (**Fig. 1F**). Confocal microscopy of the cells reflected this range of effects as well, showing distinct aggregation properties of the individual mutants (**Fig. 1G**). Based on the mCherry/GFP ratio, we grouped mutations into folded (f-Myo), and misfolded (mf-Myo) myosin, and an intermediate class (i-Myo) with varying amounts of folded and misfolded myosin (**Fig. 1F-G**). The f-Myo group includes HCM mutations that affect surface exposed residues located at the periphery of the motor domain (R403Q, R453C), i-Myo mutations affect residues of central parts of the motor domain implicated in ATPase activity (I457T)^19^, formation of the SRX state (D382Y)^19^ or UNC-45-binding (G584R)^9^, while most mf-Myo mutations affect conserved residues in the inner core of the motor domain (V606M), but also in the UNC-45-binding site (E482K, H581R) (**Fig. S2A/D**). In conclusion, data obtained with our folding reporter indicates that HCM-linked myosin mutations impair myosin folding to various degrees in cells.

### HCM mutations induce temperature-sensitive muscle defects in *C. elegans*

To investigate the effect of myosin misfolding in vivo, we introduced myopathy-linked mutations into *C. elegans*, an established model for studies of proteotoxic stress, protein misfolding diseases and age-related disorders^23-25^. Aside from using a conserved set of proteostasis factors for maintaining muscle proteins^26^, an important asset of *C. el ans* is that MHC-B myosin (encoded by the *unc-54* gene) is crucial for motility but not for viability and reproduction^27^, allowing us to characterize mutations with severe phenotypes in fully developed worms. We thus introduced HCM mutations of the f-Myo group (R403Q, R453C), the i-Myo group (G584R) and the mf-Myo group (V606M) into the endogenous mhc-B locus (**Fig. 2A**). Notably, the selected mutations are among the most common disease alleles, present in 14% of HCM patients with defective cardiac myosin^28^. As controls to selectively monitor motility defects caused by a misfolded protein, we generated a *C. elegans* strain harbouring the G463A MHC-B mutation, impairing myosin’s ATP binding and motor function^29^, and a strain with an *unc-54(0)* null allele, leading to almost complete loss of *unc-54* mRNA (**Fig. S3A**).

**Figure 2:**
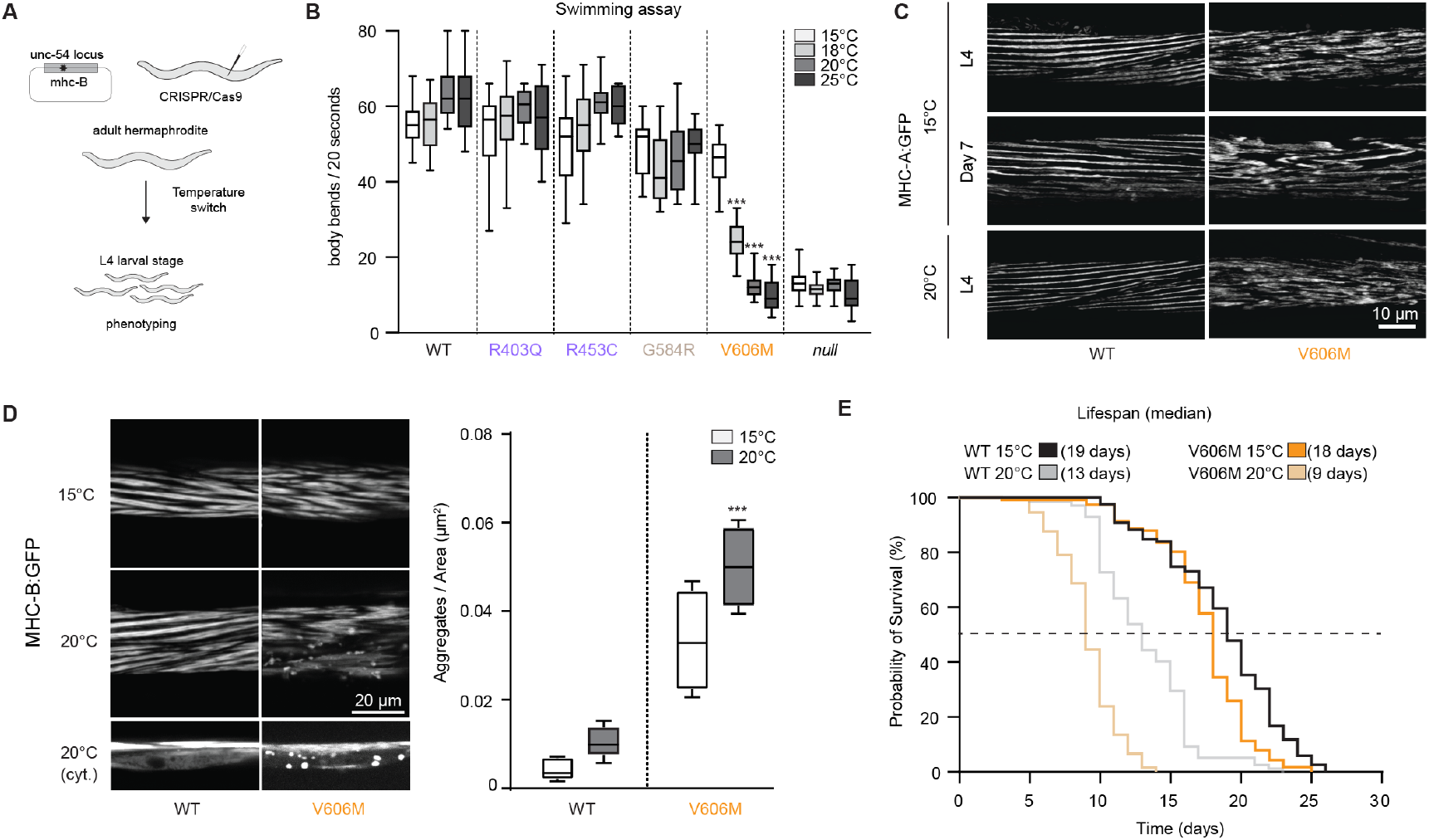
HCM mutations induce temperature-sensitive muscle defects in *C. elegans*. **A)** Schematic of *C. elegans* strain generation and experimental setup. **B)** Swimming assays of f-Myo, i-Myo and mf-Myo variants performed at increasing temperatures. For comparison, MHC-B knockout strain *unc-54(0)* is shown. **C)** Confocal microscopy of *C. elegans* body wall muscle using GFP labelled MHC-A myosin, showing the deleterious effect of folding impaired HCM variants on myofilament arrangement. Temperatures and developmental stages are indicated. **D)** Left: Confocal microscopy of *C. elegans* (L4 stage) body wall muscle, using GFP-labelled MHC-B. Images show wildtype animals and worms expressing V606M myosin at the indicated temperatures. The lower panel contains images with the focal plane being predominantly cytosolic. Right: quantification of the number of myosin aggregates formed in the indicated strains. **E)** Reduction of lifespan in V606M strains at different temperatures.

We first tested the effects on *C. elegans* muscle function by conducting motility assays at the L4 larval stage, which is a uniform and readily discernible reference point. Wildtype and mutant animals, grown at various temperatures until the L4 stage, were suspended in solution, allowing to quantify body bends in a swimming assay (**Fig. 2B**). Mutations that caused no or mild myosin folding defects (R403Q, R453C, G584R) were mostly indistinguishable from wildtype. The control mutation G463A and *unc-54(0)* allele compromised movement at all temperatures (**Fig. S3B**). In strong contrast, the V606M variant, which results in severe folding defects in the fluorescent reporter assay, caused a temperature-sensitive movement phenotype. The V606M animals were moderately impaired in motility at 15°C, which was strongly exacerbated at 18°C and 20°C, leading to near-complete paralysis at 25°C (**Fig. 2B**). This phenotype is similar to the well characterized temperature-sensitive myosin alleles *e1157* and *e1301* (**Fig. S3B**), that after sequencing revealed the mutations L352F and G389R, respectively (**Fig. S3C**, MYH7 numbering). These MHC-B variants are considered to be prone to misfolding and aggregation^30-32^, as now directly confirmed by our folding reporter (**Fig. S3D**). The G389R mutation of MHC-B is a pathology mimic of the G389E HCM mutation in human cardiac myosin^33^, making the *e1301* strain of *C. elegans* de facto a model of this myopathy.

To investigate the effect of disease mutations on muscle integrity, we crossed the myosin mutant strains with worms expressing an MHC-A:GFP fusion protein, a minor myosin isoform routinely used to visualize sarcomere organization^34^. While animals with f-Myo and i-Myo MHC-B variants had well-ordered muscle sarcomeres at all tested temperatures (**Fig. S3E**), the V606M mutation caused a temperature-sensitive distortion of myofilaments, exhibiting moderate defects at 15°C that markedly worsened at higher temperatures (**Fig. 2C**). Likewise, sarcomere defects became more severe with age, when comparing myofilaments of animals at the L4 stage and in late adulthood (**Fig. 2C, S3E**). To test the effect of disease mutations on MHC-B myosin itself, we introduced the V606M mutation in a strain expressing the endogenous MHC-B with a C-terminal GFP^35^. When worms were grown at elevated temperature of 20°C, expression of the folding-deficient V606M variant led to a disruption of MHC-B filaments and, in addition, to the formation of myosin aggregates (**Fig. 2D**). MHC-B aggregates formed predominantly in the cytosol of muscle cells rather than in sarcomeres and they were most numerous at elevated temperature (**Fig. 2D**). Over the course of 30 minutes, they did not recover from photobleaching (**Fig. S3F**), suggesting that myosin foci are stable and persistent aggregates rather than being composed of freely diffusing proteins.

Myofilament distortion and protein aggregation had also a strong impact on organismal fitness as assessed by the analysis of reproduction and lifespan of V606M animals. Specifically, at elevated temperature, the V606M strain showed an increase of bagging, a defect in egg laying frequently associated with compromised health (**Fig. S3G**), and a reduced lifespan, lowered from 23 days in wildtype worms to 14 days in mutant animals (**Fig. 2E**). Taken together, these data show that HCM-linked myosin mutations, which impair myosin folding, cause severe muscle defects in *C. elegans* at higher temperatures and with age. The misfolding of muscle myosin leads to myofilament disruption and aggregate formation, correlating with motility defects and a shorter lifespan.

### V606M destabilizes the U50 hydrophobic core of the motor domain

Temperature-dependent misfolding might either occur during myosin biogenesis or affect mature proteins incorporated into thick filaments. To distinguish between these possibilities in worms expressing V606M MHC-B, we shifted adult animals grown at 15°C to the restrictive temperature of 25°C. Unlike worms exposed to the restrictive temperature during larval development, which were paralyzed (**Fig. 2B**), the shift to a higher temperature upon completion of development resulted in a reduced decline in muscle function (**Fig. S3A**). These data suggest that HCM mutations may affect myosin synthesis rather than destabilizing the native muscle protein. To confirm this, we grew worms to larval stage L3 at 15°C, inhibited translation using cycloheximide and shifted the worms to 25°C (**Fig. 3A**). Since new myosin can’t be incorporated into the myofilaments, any phenotype should reflect the unfolding of mature myosin molecules. We observed that the motility of untreated worms declined rapidly at 25°C, whereas worms incubated with cycloheximide remained more motile in the L4 stage (**Fig. 3A**). These data show that the V606M mutation hinders the folding and assembly of myosin, rather than destabilizing myosin molecules already assembled in thick filaments.

**Figure 3:**
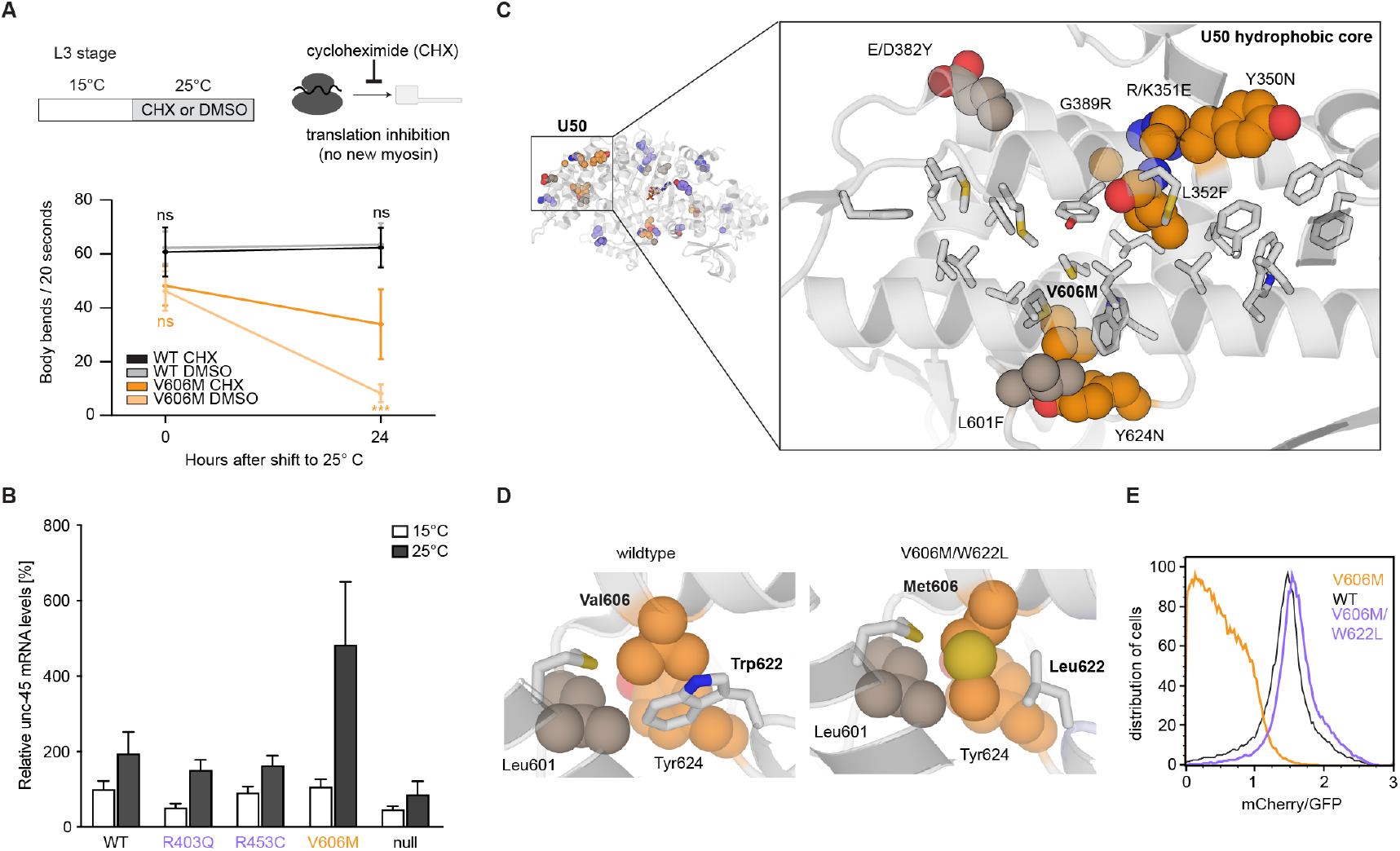
V606M destabilizes the U50 hydrophobic core of the motor domain. **A)** Inhibition of translation with cycloheximide (CHX) to address the impact of HCM mutation on myosin folding and stability of myosin assembled in myofilaments (top). Upon a temperature switch to the restrictive temperature, CHX treatment specifically improved motility of V606M worms (bottom). **B)** Analysis of mRNA levels of endogenous *unc-45* in worms expressing HCM myosin variants at permissive (15ºC) and restrictive (25ºC) temperature. **C)** Zoom-in of myosin motor domain showing that the majority of mf-Myo HCM mutations (orange) cluster at the outer rim of the hydrophobic core of the U50 scaffold (upper 50 kDa sub-domain of myosin motor). HCM mutations are shown as spheres, other U50 residues forming the U50 core in stick mode. **D)** Structural details of the neighbourhood of V606: this residue is in close contact to L601, W622 and Y624, highlighting the steric constraints of inserting a methionine at this position (left panel). Replacing the adjacent tryptophane by leucine provides space to accommodate methionine, as shown by the modelled V606M/W622L mutant (right panel). **E)** Flow-cytometry analysis of the double mutant V606M/W622L (lilac) in comparison to V606M (orange).

We therefore tested to what extent the misfolding of myosin can be mitigated by folding factors that and act in a co-translational manner. In case of myosin, such function has been reported for *Rng3*, the yeast ortholog of *unc-45*^36^. To test for a potential upregulation of *unc-45*, we determined its mRNA levels in worms expressing myosin mutants with distinct folding defects at 15°C and at 25°C. In all analysed strains, the non-permissive temperature elevated *unc-45* expression. Importantly, however, we found that the mf-

Myo mutant V606M most strongly upregulated *unc-45* expression, with mRNA levels being 4-fold increased over V606M animals grown at 15°C (**Fig. 3B**). This pronounced upregulation was not observed for other mutants, or the wildtype protein, thus UNC-45 overexpression seems to be specifically enhanced by the presence of misfolded myosin. To explore the in vivo effect of a compromised UNC-45 chaperone function, we used the *unc-45(ts)* alleles *e286* and *m9437*, which have been shown to perturb the interaction with myosin in vivo and in vitro^21,38^. When the motility of mutant worms was tested in swimming assays, the strains with defective UNC-45 exhibited temperature-sensitive motility defects similar to the V606M variant (**Fig. S4B**), connected with the appearance of myosin aggregates in body wall muscles (**Fig. S4C**). To backup these in vivo data, we tested the UNC-45(ts) variants in the split-FP reporter assay against wildtype myosin. For both variants, we observed a drastically reduced mCherry/GFP signal (**Fig. S4D**), confirming their defect in chaperoning myosin. To further link chaperone dysfunction and myosin misfolding, we tested an *unc-45* linked myopathy mutation (R754Q) located in the substrate binding domain of the chaperone. As for the *unc-45* and myosin temperature-sensitive alleles, we observed a reduced mCherry/GFP signal (**Fig. S4E**). In conclusion, our in vivo and in cell data indicate that V606M cause a defect in myosin folding that is phenocopied by a defective chaperone UNC-45.

To understand how V606M could impair myosin folding, we analysed the distribution of the folding-deficient mutations in the MHC-B structure. Strikingly, V606 is in close proximity to residues L352 and G389 mutated in the myosin ts alleles and to residues Y350, K351, D382 and Y624 implicated in HCM and causing severe protein misfolding. These residues, located far apart in primary sequence, form the outer rim of the hydrophobic core of the myosin U50 sub-domain (**Fig. 3C**), highlighting the impact of the composite U50 core for myosin maturation. Insertion of the bulkier Met (V606M) or Phe (L352F) is expected to disrupt the hydrophobic core and thus destabilize the motor domain (**Fig. 3D**). To test this prediction, we introduced compensatory mutations in the U50 core, inserting smaller residues next to HCM-linked mutations. To compensate for the V606M modification, we mutated the adjacent tryptophane W622 to leucine (**Fig. 3D**) and tested the V606M/W622L double mutant in our cellular folding assay. Strikingly, as reflected by the increased mCherry/GFP signal, introduction of W622L rescued the folding defect of V606M, yielding a mutant that has wildtype like folding properties (**Fig. 3E**). The suppressor W622L variant provides compelling evidence that the V606M mutation destabilizes myosin by disrupting the hydrophobic core of the U50 domain, yielding a muscle protein prone to misfolding and aggregation.

### HCM mutations induce severe proteotoxic stress in muscle cells

To examine the proteotoxic effect of myosin disease variants exhibiting graded folding defects, we characterized their phenotype in a chaperone-compromised environment. For this purpose, we generated double mutants by introducing HCM mutations in the *unc-45(ts) m94* background. We then assessed the motility of worms co-expressing f-Myo, i-Myo or mf-Myo mutants with the defective UNC-45 chaperone, looking for potential synthetic effects on muscle function. At the permissive temperature, the motility of worms expressing defective UNC-45 and f-Myo R403Q was only slightly compromised when compared to wildtype. In contrast, the motility of double mutants expressing f-Myo R453C or i-Myo G584R myosin in the *unc-45(ts)* background was drastically reduced (**Fig. 4A**). These data suggest that some destabilized MHC-B variants, which have no or little effect on muscle function in isolation, can induce severe muscle defects when expressed in animals with a compromised chaperone machinery. Ultimately, we attempted to generate double mutant strains bearing the *unc-45(ts)* allele together with one of the mf-Myo mutants (V606M, L352F, G389R). However, homozygous *mf-Myo*/*unc-45(ts)* double mutations were lethal at early larval stages, even at permissive temperatures (**Fig. 4A, Fig. S5A**), at which the isolated mutants otherwise display only a mild motility defect (**Fig. 2B, Fig. S3B**). In contrast to worms expressing mf-Myo, introducing the *unc-45(ts)* allele in the MHC-B null background resulted in paralyzed, yet viable animals (**Fig. 4A**). These data show that misfolded MHC-B proteins are more detrimental to muscle cells than the complete loss of myosin function.

**Figure 4:**
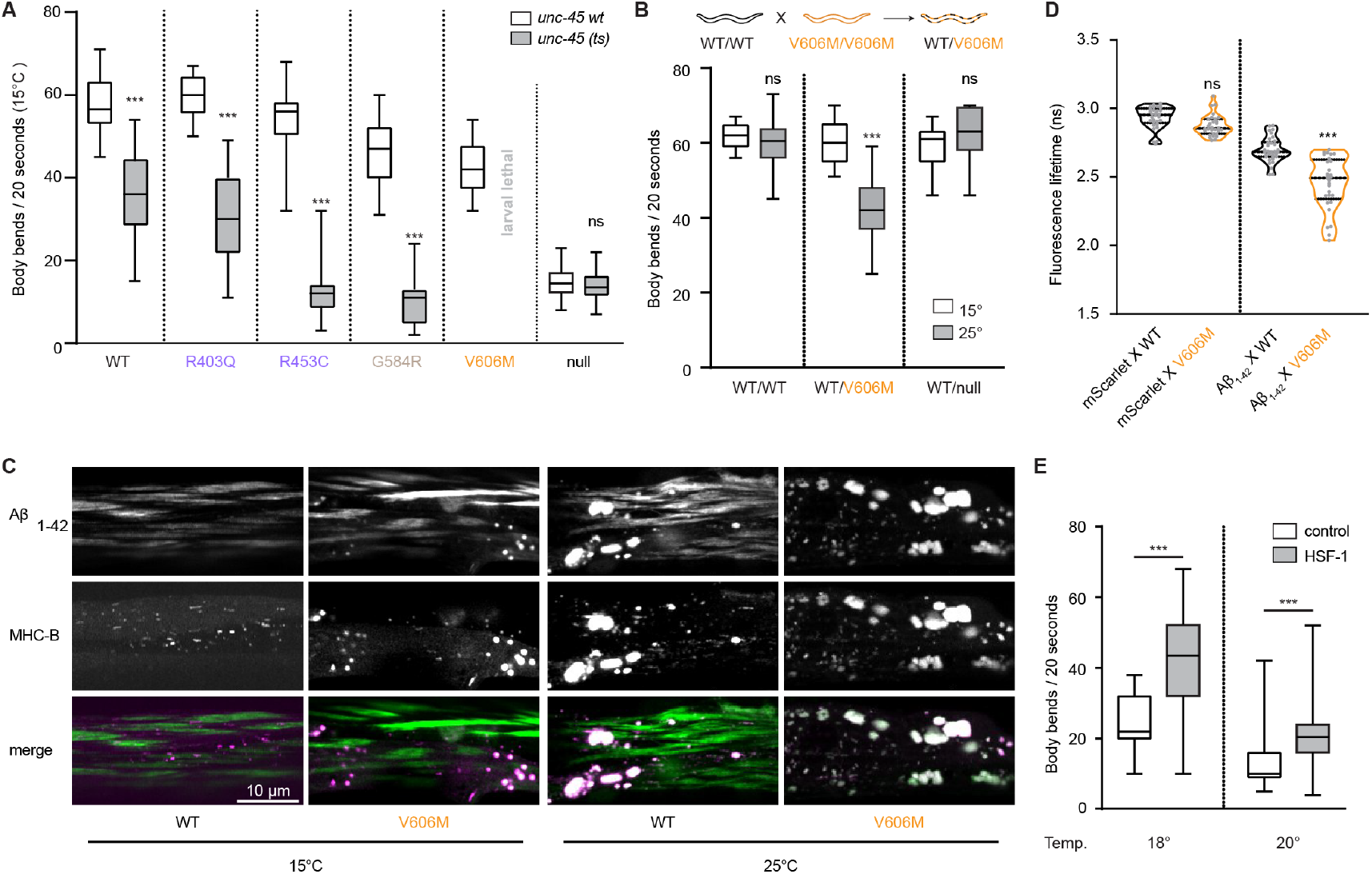
HCM mutations induce severe proteotoxic stress in muscle cells. **A)** Swimming assay monitoring synthetic effects of pathologic myosin variants and inactive UNC-45 chaperone. For the latter, we used the *m94 unc-45(ts)* strain, expressing a chaperone variant impaired in binding myosin. Contrary to catalytically defective motor proteins, mf-Myo variants showed synthetic lethality in the *unc-45(ts)* background, whereas f-Myo R452C and i-Myo G584R were strongly impaired in motility. **B)** Crossing of wildtype (WT) and V606M worms yielded WT/V606M heterozygous worms that showed a significant reduction in motility at high temperatures, contrary WT/null animals were not impaired in motility. **C)** Confocal microscopy of *C. elegans* (day-4 old animals) body wall muscle, using fluorescently labelled Aβ^1-42^ (mScarlet) and MHC-B (GFP) variants. Experiments were performed in WT and V606M animals, at 15°C and 25°C. **D)** Fluorescence lifetime of mScarlet and mScarlet-Aβ^1-42^ in wildtype (WT) and V606M myosin background. **E)** Rescue of the motility defect by overexpression of the HSF-1 transcription factor.

To explore the toxic effect of myosin misfolding, we generated heterozygous worms expressing both wildtype and V606M myosin. To confirm the equal expression level of the alleles in the heterozygous background, we performed allele-specific qPCR against wildtype *unc-54* and *unc-54(V606M)* mRNA. The two alleles were equally expressed (**Fig. S5A**), ruling out an effect of the mutation on transcript stability in *C. elegans*, as previously proposed for the V606M mutation in cardiac myosin^39^. In the motility assay performed at 15°C, the heterozygous animals behaved like wildtype. At 25°C, however, the motility of wildtype/V606M worms was strongly impaired (**Fig. 4B**). Moreover, lack of one mhc-B allele in heterozygous wildtype/null animals had no negative impact on motility (**Fig. 4B**), indicating that lower levels of MHC-B do not impact muscle function. Taken together, these data indicate that misfolded MHC-B myosin has a proteotoxic effect in muscles.

Diverse protein aggregates can act synergistically, resulting in a worsening of the phenotype as shown for poly-glutamine Q40 aggregates in paramyosin ts strains^31^. To test for aggregates synergistic proteotoxicity, we crossed V606M worms with nematodes expressing the aggregation-prone model substrate Aβ^1-42^ in muscle cells^40^. We used Aβ^1-42^ amyloid aggregates as a molecular reporter of proteotoxicity as well as an additional stressor elevating the proteotoxic burden in muscle cells. In contrast to the parental V606M strain, expression of Aβ^1-42^ induced aggregation of V606M myosin even at the permissive temperature (15ºC), highlighting the latent aggregation propensity of this HCM mutant (**Fig. 4C**). Likewise, the presence of V606M myosin enhanced the aggregation of Aβ^1-42^, reflected by a reduction of fluorescence lifetime of mScarlet fused to Aβ^1-42^, (**Fig. 4D**).

At the restrictive temperature of 25ºC, the proteotoxic stress was more severe. Wildtype myosin started aggregating (**Fig. 4C**), a phenotype similar to what we observed in worm with defective UNC-45 chaperoning (**Fig. S4D**). In the mutant myosin background, we observed a further increase in protein aggregation of both V606M myosin and Aβ^1-42^, culminating in an almost complete distortion of MHC-B filaments. This extreme phenotype, which significantly damages muscle structure, demonstrates the cumulative impact of distinct proteotoxic stresses, which can overwhelm the proteostasis machinery in muscle cells. Remarkably, in contrast to Q40/paramyosin aggregates that form separately^31^, Aβ^1-42^ co-aggregated with V606M myosin (**Fig. 4C**). This suggests that aberrant proteins can act as a seed, amplifying the aggregation tendency of misfolded myosin molecules and resulting in the formation of mixed protein aggregates.

Given the connection of disease mutations, protein aggregation and muscle dysfunction, we hypothesized that enhancing proteostasis pathways in muscle cells may relieve the ts phenotype of HCM alleles. To test this, we over-expressed in body wall muscle cells the heat shock transcription factor *hsf-1*, a central regulator of the stress response also controlling chaperone expression. *hsf-1* overexpression in body wall muscle resulted in a partial rescue of the motility defect at 18°C and 20°C (**Fig. 4E**), indicating that boosting the heat shock response can mitigate the harmful effect of myosin misfolding. Taken together, our data demonstrate that misfolded myosin has a dominant-negative effect on the proteostasis system in muscle cells, and that these deleterious effects can be enhanced by additional proteotoxic stresses, such as the presence of other aggregation-prone proteins.

### Caloric restrictions can mitigate HCM defects via autophagy

Dietary restriction is highly effective in stimulating the integrative stress response that counteracts protein aggregation^41,42^. Moreover, caloric restriction protects against sarcopenia, the loss of muscle proteins with age^43^, suggesting that it might mitigate muscle defects caused by HCM-linked myosin misfolding. To test this hypothesis, we compared the motility of well-fed and starved worms, expressing HCM-linked myosin variants at the restrictive temperature. We found that food withdrawal resulted in a remarkable four-fold increase in motility for V606M worms as compared to well-fed animals (**Fig. 5A**) and observed a similar, persistent rescue with other mf-Myo strains (**Fig. S6A, S6B**). To a minor extent, the rescue is due to a behavioural component as food withdrawal from wildtype, R403Q and R453C animals also influenced motility (**Fig. 5A**). Nevertheless, the motility of worms expressing the inactive variant G463A or the myosin null allele was not recovered by starvation (**Fig. 5A, S6A**). Together, these findings show that caloric restriction induces a specific stress response to counteract proteotoxic stress by misfolded myosin.

**Figure 5:**
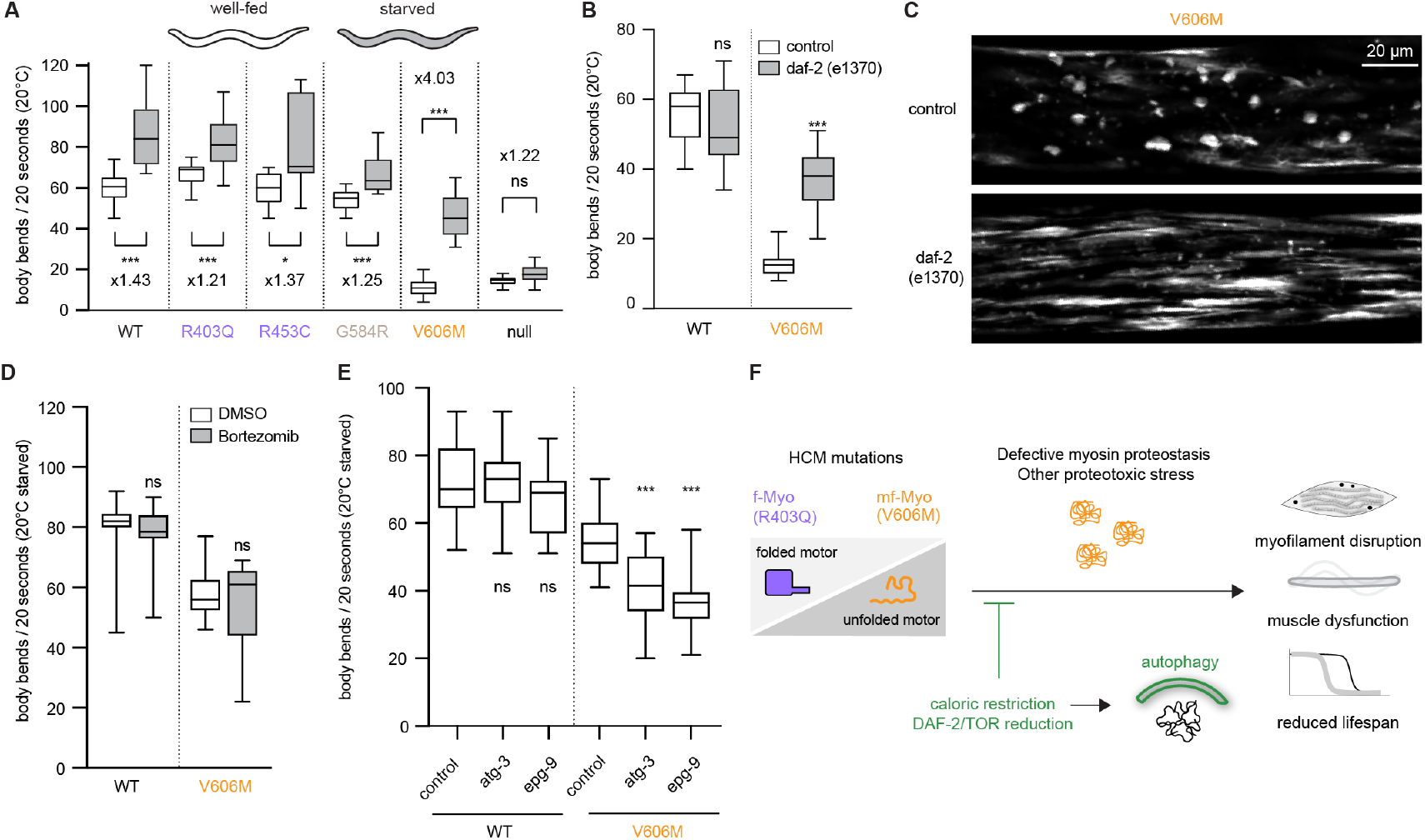
Caloric restrictions can mitigate HCM defects via autophagy. **A)** Swimming assay comparing motility of well-fed and starved worms at the L4 stage. The tested myosin variants are indicated. Specifically, for V606M we observed a 4-fold increase in motility. **B)** Inhibition of *daf-2* signaling led to pronounced rescue in motility, specifically in worms expressing folding deficient myosin. **C)** Confocal microscopy of WT and *daf-2* mutant (*e1370*) worms at their L4 stage, using GFP-labelled MHC-B. Inhibition of *daf-2* signaling mostly reduced aggregats, restoring the muscle structure. **D)** Effects of proteasome inhibition on the starvation rescue. **E)** Effect of autophagy reduction on the starvation rescue. Contrary to proteasome inhibition, a reduction of the rescue is observed. **F)** Model of myosinopathies: myosin aggregation in muscle can be caused by point mutation, and be exacerbated by defective proteostasis and other stresses, resulting in both a loss of function and a toxic side effects on cellular and organismal levels. Starvation and inhibition of *daf-2* and TOR pathways result in a rescue of the phenotype, mediated by autophagy.

Insulin and TOR signalling are among the major pathways linked to nutrient sensing. We thus tested the contribution of TOR signaling for improving muscle function in V606M worms by generating double mutants with variants in *rict-1*, a component of the TOR signaling complex inhibited during starvation. Introduction of the *rict-1* alleles *mg360* (a missense mutation)^44^ or *ft7* (a nonsense mutation)^45^ significantly enhanced motility of V606M worms at the restrictive temperature (**Fig. S6C**). To confirm the rescue by TOR signaling, we depleted by RNAi the TOR kinase, encoded in *C. elegans* by the *let-363* gene. The motility of worms expressing folding-deficient MHC-B mutants was significantly improved upon LET-363 depletion, whereas worms expressing inactive MHC-B mutants or the myosin null allele were unaffected (**Fig. S6D**). To test the contribution of insulin signalling on myosin proteostasis, we crossed the V606M worms to the *daf-2* allele *e1370*, a partial loss-of-function variant of the insulin-like growth factor 1 receptor^46^. Strikingly, the muscle function of V606M worms was almost fully restored in the *daf-2(e1370)* background at the restrictive temperature (**Fig. 5B**), phenocopying the rescue effect by starvation. We next investigated to what extent inhibition of *daf-2* could prevent damage in the ultrastructure of muscle cells. Confocal microscopy of worms expressing labelled MHC-B showed that *daf-2* inhibition strongly reduced myosin aggregates in number (**Fig. S6E**) and size (**Fig. S6F**), while at the same time improving the alignment of myofilaments (**Fig. 5C**).

Insulin signalling is known to regulate the ubiquitin-proteasome system by inhibiting its key regulator, the transcription factor *skn-1* (skinhead-1)^47^. To determine if the UPS pathway plays a role in mitigating proteotoxic stress caused by aggregated myosin, we assessed the motility of starved V606M worms in both a *skn-1a* deletion background (**Fig. S6G**), or by applying the proteasome inhibitor bortezomib (**Fig. 5D**). Neither condition abolished the rescue in motility, suggesting that the UPS pathway does not significantly contribute to counteracting myosin misfolding and aggregation under these conditions. In addition to the UPS, insulin signalling upregulates autophagy pathways in muscle. To investigate whether autophagy is important for the rescue of motility upon starvation, we exploited the autophagy hypomorph strains *epg-9* and *atg-3* and crossed them to the V606M strain.

The autophagy alleles did not impact movement in the wildtype background during starvation (**Fig. 5E**). However, in worms expressing V606M MHC-B, we observed a significantly reduced rescue upon starvation (**Fig. 5E**). Silencing of *hlh-30*, which encodes the *C. elegans* homologue of the autophagy master regulator TFEB, also resulted in a similar reduction of starvation-induced motility rescue (**Fig. S6H**). Taken together, these findings demonstrate that the inhibition of insulin/TOR signaling upon starvation initiates an autophagy-driven proteostasis response that mitigate defects in muscle structure and function caused by misfolded and aggregated myosin.

## DISCUSSION

Protein folding and degradation pathways are essential for maintaining cellular homeostasis, and their failure is a hallmark of aging and various diseases. In this study, we developed a split-FP reporter to systematically assess the folding capacity of pathological myosin variants linked to cardiomyopathies. Our results reveal that HCM mutations in the myosin motor domain can induce protein misfolding, defining a distinct class of myopathy mutations. Upon establishing a *C. elegans* myopathy model, we demonstrated that HCM-linked folding defects lead to myosin aggregation, sarcomere disruption, and compromised muscle function, effects that can be mitigated by dietary restriction and inhibition of insulin and mTOR signaling pathways.

Assessing protein folding in cells is technically challenging, as existing fluorescent protein tags typically report on protein distribution and aggregation but do not directly indicate the folding state. To address this gap, we incorporated a split-FP into the myosin scaffold such that it only emits a signal once the motor protein adopts its folded state. Using this reporter in a flow-cytometry based setup, we revealed that disease mutations impair myosin folding to varying extents. These findings align with human data showing distinct abundances of severely misfolded (V606M) and mildly misfolded (G584R) alleles in patients^39^, highlighting the potential of our split-FP as a diagnostic tool for classifying HCM mutations. Importantly, our approach identified protein variants with minor folding defects that have remained undetected in traditional assays, suggesting that many HCM mutations result in metastable proteins with latent pathologic potential. Consistent with this, a recent study showed that metastable proteins can maintain a pseudo-wildtype phenotype while inducing broad proteomic alterations under stress conditions^48^. In our *C. elegans* myopathy model, even subtle folding defects linked to HCM mutations were drastically exacerbated when the chaperone machinery was disrupted. Notably, missense mutations that affect protein stability can also lead to protein mislocalization, presenting a widespread disease mechanism^49^. Taken together, these data suggest that in human cardiomyocytes, folding-impaired myosin variants may be buffered by specialized chaperones and degradation factors for extended periods, with their full pathological effects only becoming apparent under conditions of proteotoxic stress or aging when the proteostasis system is overwhelmed. Supporting this notion, the V606M mutation does not yield an apparent phenotype in mouse models, unless the mice are treated with cardiotoxic drugs or are crossed with other HCM alleles^50^. We thus propose that in analogy to the here characterized myopathies, a substantial number of missense mutations in the human proteome should result in metastable proteins, with their full pathological potential only becoming apparent when the cellular proteostasis network is overwhelmed. Likely candidates for such latent protein disorders include other muscle proteins, like titin, as well as neuronal isoforms of tubulin^51-53^.

Our study indicates that many HCM-linked myosin mutations cause folding defects rather than disrupt ATPase activity. The identified folding-deficient mutants cluster in two key regions, the binding site for the UNC-45 chaperone and the hydrophobic core of the U50 sub-domain. Notably, an unbiased screen for mutations yielding metastable proteins and upregulating proteasomal expression in *C. elegans* identified mutations in myosin U50 as well^32^. This suggests that U50 assembly poses a significant challenge to co-translational folding, requiring the structural alignment of distant protein regions while accommodating the UNC-45 chaperone. Given the apparent folding constraints, formation of the U50 sub-domain may act as a regulatory checkpoint, relying on translation dynamics, codon preferences, and auxiliary folding factors to ensure efficient myosin maturation. Interestingly, the deleterious effects of disease mutations in this region can be rescued by compensatory mutations in adjacent residues, suggesting that pharmacological chaperones can be developed to target metastable myosin variants - a concept successfully applied in cystic fibrosis to restore protein function^54^.

As HCM mutations impact either myosin activity or folding, they necessitate distinct therapeutic approaches. For mutations that deregulate ATPase activity, current treatments involve small molecules that stabilize the super-relaxed SRX state and reduce ATP consumption^55,56^. Conversely, mutations like V606M that induce misfolding and aggregation will require alternative strategies, such as small-molecule modulators of proteostasis pathways. Our data demonstrate that dietary restriction and inhibition of insulin/TOR signalling not only reduce myosin aggregation but also improve muscle function in *C. elega*. Interestingly, the positive effects of caloric restriction and TOR inhibition on hypertrophy have been shown before^57,58^, but not for myosin-based myopathies. In worm expressing misfolded, aggregation-prone myosin, the rescue effect was strongly dependent on autophagy, as mutations in *epg-9* and *atg-3* impaired the benefits of starvation. Consistent with this, FOXO1, a downstream effector of insulin signalling critical for autophagy upregulation^59^, is implicated in the regression of cardiac hypertrophy in vertebrates^60^. In conclusion, our findings underscore the central role of autophagy in maintaining muscle cell proteostasis, particularly in response to proteotoxic stress induced by misfolded myosin. Enhancing autophagic pathways therefore presents a compelling therapeutic strategy for myosin based HCM and other protein misfolding disorders in muscle cells.

## ACKNOWLEDGMENTS

We thank Luisa Cochella for sharing strains and reagents. We acknowledge the imaging core facility of the FLI for their support and thank Jana Neuhold from the VBCF Protein Technologies facility for support in protein engineering as well as all members of the Clausen and Kirstein labs for discussions.

## AUTHOR CONTRIBUTIONS

Conceptualization: RA, PJD, TC, JK Methodology: RA, PJD, TC, AM, JK

Investigation: RA, PJD, BED, LK, GA, NK, JK, LD Visualization: RA, PJD, TC

Funding acquisition: RA, TC, JK Project administration: RA, TC, JK Supervision: TC, JK

Writing – original draft: RA, PJD, TC Writing – review & editing: RA, TC

## DATA AND MATERIAL AVAILABILITY

All data are available in the main text or the supplementary materials. C.elegans strains will be available upon request

## MATERIAL AND METHODS

### Materials Availability

Cells and plasmid deposition: Important *C. elegans* strains will be made available at the Caenorhabditis Genetics Center (University of Minnesota), other strains and plasmids are available upon request.

### Design and cloning of the fluorescent reporter

All sequences were amplified by Phusion Hot Start Flex II polymerase (NEB) according to manufacturer’s instructions. The backbones used in this assembly are derived from the biGBac system^2^. For the fluorescent reporter heavy chain, GFP, MHC-B and mCherry fragment 160-239 were amplified and assembled with Gibson Assembly Mix from N-term to C-term in a pLib plasmid, with a GGGSGGGS linker between MHC-B and both fluorophores. For the fluorescent reporter light chain, MLC-3 and mCherry fragment 1-159 were amplified and assembled with in-house Gibson Assembly Mix from N-term to C-term in a pLib plasmid, with a GGGSGGGS linker between MLC-3 and mCherry. UNC-45 was amplified and assembled with Gibson Assembly Mix a pLib plasmid. The final assembly containing MHC-B, MLC-3 and UNC-45, in the accepting backbone pBIG1a was generated as reported previously^62^. To generate point mutations of the folding reporter of myosin or UNC-45, mutagenesis with PCR was performed.

### Viral production

Plasmids expressing the expression vectors and fluorescent reporters were transformed in DH10EMBacY or DH10Bac cells, respectively. The second cells don’t have any fluorescent marker of infection to not interfere with the detection of the GFP signal. After blue/white screening, bacmids were extracted by alkaline lysis and isopropanol precipitation. Baculoviruses were produced transfecting Sf9 cells (Expression Systems) in adhesion in 6-well plates (1 million cells in 3 ml of ESF-921 medium) with the bacmid (3 μg) and PEI (10 μl, 1 mg/ ml concentration). 7 days after transfection, Sf9 insect cells in suspension culture were infected with supernatant containing the virus from the previous step at 27°C and 100 rpm for 4 days. The V1 generation was harvested, filtered, and stored at 4°C.

### Flow-cytometry analysis of fluorescent reporter

Hi5 cells (Expression Systems) were grown to a density of 1×106 cells/mL in suspension and infected with 1:50 dilution of virus. After incubation for 4 days at 21°C and 100 RPM, cells were filtered through filter-cap tubes for flow-cytometry analysis with LSR Fortessa. Control cells infected with UNC-45 only were used to set a threshold for positivity of both GFP and mCherry signals. Positive cells were gated and the ratio of mCherry/GFP calculated for at least 30000 cells per experiment. Data plotting and analysis was performed with FlowJo.

### Solubility assay in insect cells

To correlate the mCherry/GFP ratio to the solubility of the fluorescent reporter, cells were pelleted at 1000G at 4°C for 20 minutes, weighted and flash-frozen in liquid nitrogen. The pellets were then resuspended in the following lysis buffer: 50 mM Na phosphate pH 8, 300 mM NaCl, 1:5000 benzonase and 1 tablet of cOmplete protease inhibitor (Roche) every 50 ml of buffer. The volume of buffer for the resuspension was proportional to the weight of the pellet (5X). After resuspending the cell pellet, 500 μl of each sample were centrifuged at 20000G for 30 minutes at 4°C to separate soluble and insoluble proteins. The supernatant (Sup) was separated from the pellet, which was resuspended in the same amount of buffer, and both were loaded on SDS-PAGE.

### Protein expression and purification

WT, V606M and V606M/W622L myosin variants were expressed and purified as previously reported^9^. Briefly, the MHC-B variants were cloned in a pFastBac DUAL vector together with UNC-45, and baculoviruses were produced (see Viral Production). 1 liter of Hi5 cells (Expression Systems) at a density of 1 × 106 cells ml^−1^ were infected with the virus and incubated 4 days at 21°C. Cells were pelleted and flash frozen in liquid nitrogen, in 50 mM Tris, pH 7.5, 300 mM NaCl (buffer A), and one tablet of cOmplete protease inhibitor (Roche) every 50 ml of buffer. After centrifugation at 20000G for 30 minutes at 4°C, the supernatant was loaded onto a NiNTA column (GE Healthcare), and after washes with increasing imidazole, eluted with buffer A plus 200 mM imidazole. The elution was then applied to a S200 columns (GE Healthcare) pre-equilibrated with 20 mM Tris pH 8.5, 20 mM NaCl, and eluted with the same buffer.

### Animal maintenance

*C. elegans* was maintained at 15°C in the dark on solid nematode growth media (NGM) seeded with live OP50 E. coli. All assays were performed with age-synchronized nematodes at the indicated temperatures. Animals were synchronized by egg-laying or by picking L4 larvae.

### Genetic crosses

Male nematodes used for genetic crosses were generated by heat shock of L4 larvae/young adults at 30°C for 7 h. Crosses were performed by transferring 1 L4 hermaphrodite of one genotype with 6 males of the other genotype. F1 offspring were self-fertilized to obtain homozygous F2. Correct crosses were validated by analyzing the fluorescence signals or by genotyping. To generate compound heterozygote animals, males which were homozygous for a specific *unc-54* allele and expressed *rpt-3::gfp* from an integrated transgene were mated to GFP-negative hermaphrodites homozygous for the complementary *unc-54* allele, and GFP-positive F1 progeny were collected for analysis. Some strains were provided by the CGC, which is funded by the NIH Research Infrastructure Programs (P40 OD010440).

### Transgenesis and genome engineering

To introduce specific mutations at the *unc-54* locus, animals were subjected to CRISPR/Cas-9 mediated genome editing via microinjection of RNPs as described by Paix et al., 2014, with modifications from Prior et al., 2017. We employed the Alt-R CRISPR/Cas9 system (Integrated DNA Technologies), animals were injected with a mixture containing 300mM KCl, 20mM HEPES, 4mg/ml recombinant Cas9 (from *S. aureus*, purified in-house), 2.5ng/µl pCFJ90 (encoding the *myo-2p::mCherry* selection marker), 500ng/µl TracerRNA, 100ng/µl crRNA targeting the editing site, and 50ng/ul single stranded donor oligonucleotide serving as a repair template. Successful editing events were identified by Sanger sequencing, for sequences of crRNAs used and alleles generated see supplementary table S2. Animals bearing an extra-chromosomal array which expresses *hsf-1 c*DNA under the control of the *unc-54* promoter were generated via microinjection^63^. The injection mix contained 100ng/ µl sonicated *E*.*coli* DNA, 1ng/µl linearized pUC19 vector encoding *unc-54p::hsf-1(cDNA)::tbb-2*, and 1 ng/ µl pCFJ90 encoding *myo-2p::Cherry* as a co-injection marker.

### Bulk worm synchronization

To obtain a population synchronized in age, 5 NGM plates were filled with 5-7 worms at L4 stage. Plates were incubated at 15° C until a population with mostly gravid adult worms was observed (usually after 5-7 days). Worms were washed from the plates with M9 buffer and collected in a 15 ml centrifugation tube by a 3 min spin at 750 g, 4° C with the breaking ramp set to 4. To break the cuticles, after washing the worm pellet 3 times in M9 buffer, 10 ml bleaching solution (1 M NaOH, sodium hypochlorite 1:10) was added to the pellet. Heavy shaking and vortexing was performed for 3-4 min, while observing the process in between under the dissecting microscope. 5 ml of M9 buffer were added and the eggs were collected at the bottom of the tube by centrifugation (3 min, 750g, 4° C, breaking ramp 4). Three additional washes in M9 buffer were performed to remove remaining bleaching solution. Finally, the eggs were dissolved in 10 ml M9 buffer and incubated at 15° C for at least 24 h. On the next day, freshly hatched synchronized larvae were plated onto NGM plates and incubated at the desired temperature.

### Swimming assays

Functionality of body wall muscle cells was assessed in a thrashing assay by counting the frequency of lateral swimming movements animals perform while suspended in M9 buffer (22 mM KH_2_PO_4_; 43 mM Na_2_HPO_4_; 86 mM NaCl; 1 mM MgSO_4_) over a period of 20 seconds. Compared to tracking animal movement on plates, thrashing assays offer a more sensitive measurement allowing to discriminate minor changes in muscle function. For experiments, videos were recorded under brightfield conditions using a Zeiss SteREO Discovery.V8 microscope equipped with a Chameleon®3 Monochrome Camera (FLIR). Videos were analysed manually in a semi-blinded fashion, e.g. the scoring individual had no knowledge of genotype and condition at the moment of analysis. One body bend was defined as a complete bend in one direction. Data for a particular individual was discarded if the animal was visibly damaged or remained still for a duration of >5 s. Unless for starvation experiments, only well-fed animals that exhibited no signs of starvation, like shallowness due to low intestinal fat-content, were selected. To control for environmental variations, locomotion rates of different strains within one experiment were assayed on the same day.

### Life span assay

Worms were first synchronized as described and incubated at 20°C. When the synchronized populations reached L4 stage, 3×40 worms per strain were transferred to a fresh NGM plate, seeded with OP50. For approximately 7 days, the initial worms were transferred to a fresh NGM plate every day, to avoid confusion with ageing progeny. The condition of every worm was observed and noted every day until the end of the life span. Worms that crawled to the wall of the plate and died of starvation or worms that died because of internal hatching of larvae were not taken into analysis. Statistical analysis was done using GraphPrism and performing a Mantel-Cox test.

### RNAi knockdown

Knockdown was induced via RNAi by feeding. To this end *E. coli* HT115 bacteria expressing dsRNA against target genes of interest were selected from the Vidal ORFeome library64. For genes that were not present in the ORFeome library, plasmids were cloned manually by inserting 500 bp of cDNA-derived mRNA coding sequence into the L4440 double T7 RNAi feeding vector prior to electroporation into *E. coli* HT115. RNAi vectors were grown in liquid culture in LB-medium supplemented with 50 μg/ml Tetracyclin. A fraction of the culture was used to isolate plasmid for verifying identity of each clone via Sanger sequencing. The rest of the culture was pelleted, resuspended in LB-medium without Tetracyclin, and seeded a 10x concentrate on standard NGM plates containing 50 μg/ml Carbenicillin as well as 1 mM IPTG. Bacteria were allowed to grow overnight at 37°C prior to transferring young adult worms onto plates and analysing F1 progeny in the L4 stage. Potency of RNAi-plates was verified via *ama-1* (RNA Pol II) knockdown-mediated larval lethality in every experiment.

### mRNA expression levels

mRNA levels were quantified via RT-qPCR. Per sample 5 L4 animals were transferred to 2.5 µl of Lysis Buffer (5 mM Tris-HCl pH 8.0, 0.25 mM EDTA and 1 mg/mL Proteinase K, 0.5% Triton X-100, 0.5% Tween-20) using a thin glass needle. Samples were subjected to 10 min digestion at 65°C before heat inactivation of proteinase K for 1 min at 85°C. Next, crude lysates subjected to reverse transcription using SuperScript III (ThermoFisher) and qPCR was performed using the GoTAQ qPCRMastermix (Promega) according to the manufacturer’s instructions. Relative expression was calculated via the ΔΔCq-method using *cdc-42* as a reference gene, the following primer sequences were employed:

unc-54-F: 5’-CTGCTATGCTCATCTACACCT-3’

unc-54-R: 5’-TGTGGTGGCATTTCTGTCTT-3’

unc-45-F: 5’-GCTGATGAATTATACACTGAAGC-3’

unc-45-R: 5’-GAGCCTCTTTTGCGTCTTGA-3’

cdc-42-F: 5’-TGGGTGCCTGAAATTTCGC-3’

cdc-42-R: 5’-CTTCTCCTGTTGTGGTGGG-3’

For quantifying allele-specific levels of *unc-54(wt)* and *unc-54(V606M)* mRNA, compound heterozygotes were collected and processed as described above before cDNA was subjected to TaqMan (Applied Biosciences) qPCR using custom designed allele-specific hydrolysis probes. Relative abundance was calculated via the ΔΔCq-method using *pmp-3* as a reference gene.

### Pharmacological treatments

For treatment of animals with cycloheximide and bortezomib, stock solutions of either compound were prepared in DMSO. Compounds were then diluted in M9-Buffer to desired concentrations and layered on top of regular NGM plates seeded with *E. coli* OP50. For the cycloheximide experiment, L3 animals were reared at 15°, collected and measured in a swimming assay before being transferred 25°. Transfer plates contained either 0.25 mg/ ml cycloheximide or 0.125% DMSO. Animals were put for 1h at 15° before being shifted to 25°C and assayed 24 h later.

### Microscopy

To visualize sarcomere structure, animals expressing *myo-3p::gfp* or GFP-tagged MHC-B were mounted onto a thin agar pad on a microscopic slide and anaesthetized using 200 mM NaN3. GFP fluorescence Images were acquired via an Olympus IX3 Series (IX83) confocal microscope equipped with a Yokogawa W1 spinning disk and a sCMOS (ORCA-Fusion) camera using the CellSens Dimension software. For the reporter of folding, Hi5 cells (Expression Systems) were grown to a density of 1×106 cells/mL in suspension and infected with 1:50 dilution of virus. After incubation for 4 days at 21°C and 100 RPM, cells were placed in a µ-Dish 35 mm high (IBIDI), and imaged. For the β-amyloid experiments, confocal images of body wall muscle were obtained using a Zeiss LSM 880 microscope in ‘airyscan’ mode under a Plan-Apochromat 63×/1.4-NA objective. Images within one experiment were collected using the same laser power and exposure time and processed identically in ImageJ.

### FRAP experiments

To test dynamics of MHC-B(V606M)::GFP aggregates, FRAP experiments were performed using an Olympus IX3 Series (IX83) confocal microscope equipped with a Yokogawa W1 spinning disk, a sCMOS (ORCA-Fusion) camera, and a FRAP unit. Animals were anaesthetized and mounted as described above, and individual aggregates were identified under 100x magnification. Aggregates were photobleached using a 405 nm FRAP Laser and relative fluorescence profiles pre-and post-bleach were quantified using ImageJ. To this end, pixel intensity along a transverse section through the aggregate in a single focal plane was measured at different timepoints and normalized to the maximum pixel intensity = 1. All images within one experiment were acquired using identical laser powers and exposure times.

### Aggregate quantification

Image analysis of muscle aggregates using machine learning feature selection was performed blindly using the ilastik open-source image classification and segmentation software (www.ilastik.org)^65,66^. Ilastik uses machine learning algorithms to automatically, but under supervision, quantify features in an image. Feature selection algorithms were set with a hysteresis method for the aggregates label at a core threshold of σ = 0.6 and a final threshold of σ = 1.0 with a minimum size of 10 pixels and not merging the objects. Five labels were used in the training modules; one to identify and train for aggregates, and one to teach the exclusion of background noise. The other three will help to identify and exclude confounding features in the images, such as muscle fibres, cytoplasm of the muscle cells and vesicles. After training, experimental images were batch processed, and object information was exported and used for the number quantification and size measurements. T-tests were undertaken to assess statistical significance.

### Fluorescence lifetime

Fluorescence lifetime imaging of mScarlet and Aβ^1-42^-mScarlet animals was carried out in a Leica SP8 confocal microscope through a Plan-Apochromat 40x/1.3-NA objective. Both acquisition and analysis were done using ‘LASx’ and ‘Single Molecule Detection’ software modules (Leica Microsystems GmbH). A pulsing white light laser at 80 MHz was used for excitation of mScarlet at 561nm. Emitted photons at 575nm - 624nm were counted by a Time Correlated Single Photon Count (TSCPC) detector ‘Leica Hyd’ at a rate below 1% of laser repetition rate until at least 1500 photons were collected from a brightest pixel. Mean fluorescence lifetime (τ) value for each acquisition was obtained by fitting the fluorescence decay curves from 2000 ns – 7000 ns in ‘mono-exponential tail fit’ mode.

### Statistical analysis

GraphPad Prism version 8 was employed for statistical analysis: normality test was performed using D’Agostino-Pearson test, while One-way ANOVA with Tukey post hoc test or t-test were used depending on the experiment. No exclusion criteria were pre-determined, and no animals or data such as outliers were excluded from subsequent statistical analyses. P-values: ns = p > 0.05; * = p ≤ 0.05; ** = p ≤ 0.01; *** = p ≤ 0.001.

## SUPPLEMENTARY FIGURES

**Figure S1:**
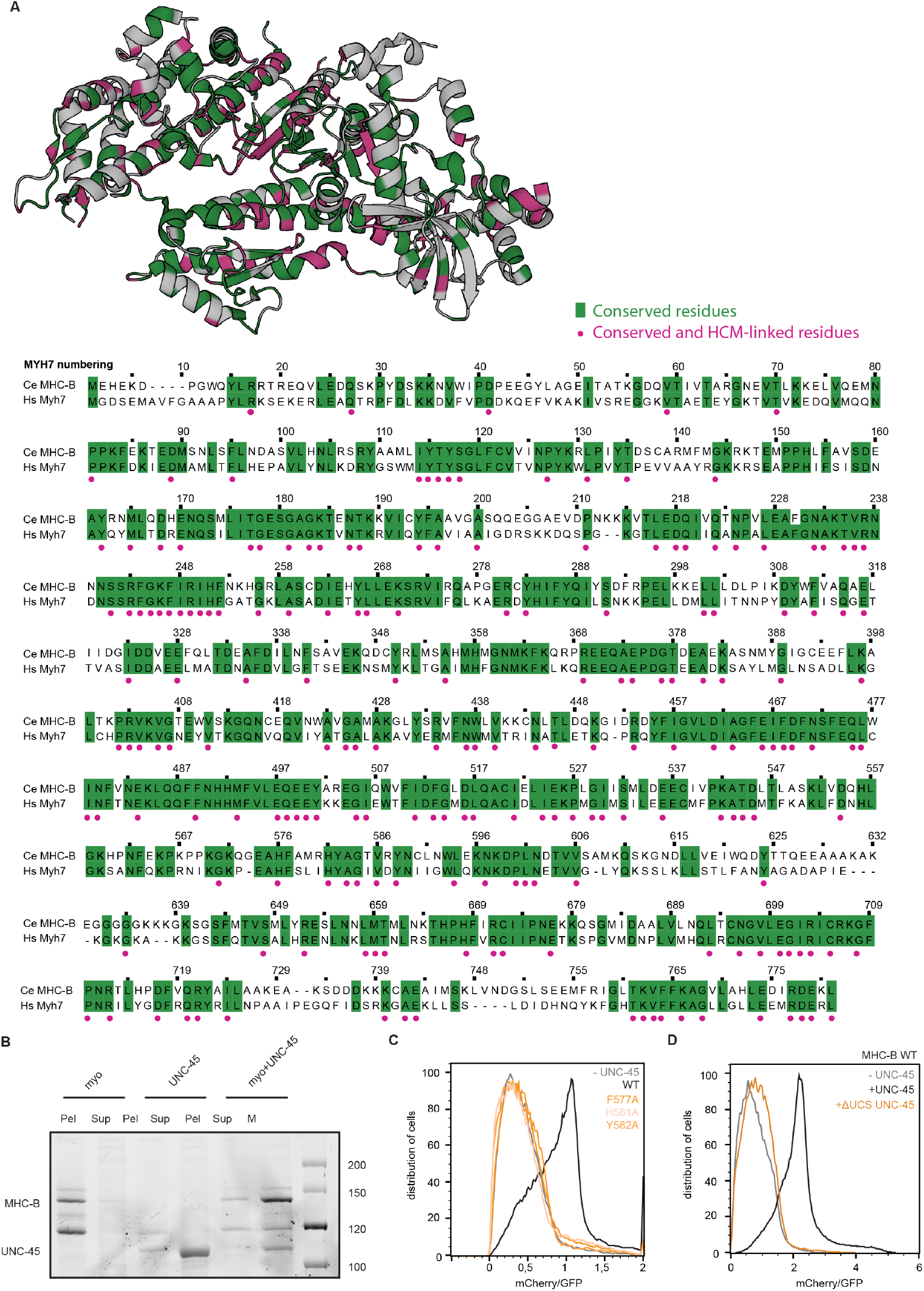
HCM mutations cause graded folding defects in myosin. **A)** Alignment of *C. elegans* MHC-B to human β-cardiac myosin. Identical (conserved) residues are highlighted in green, whereas a magenta dot represents conserved residues linked to cardiomyopathies, as listed in the Human Gene Mutation Database^61^. Both conservation and disease mutations were mapped on the structure of MHC-B. **B)** SDS-PAGE of soluble and insoluble fractions of MHC-B myosin folding reporter in presence or absence of UNC-45. **C)** Flow-cytometry analysis of MHC mutants with WT UNC-45. **D)** Flow-cytometry analysis of the ΔUCS UNC-45 mutant with WT MHC-B.

**Figure S2:**
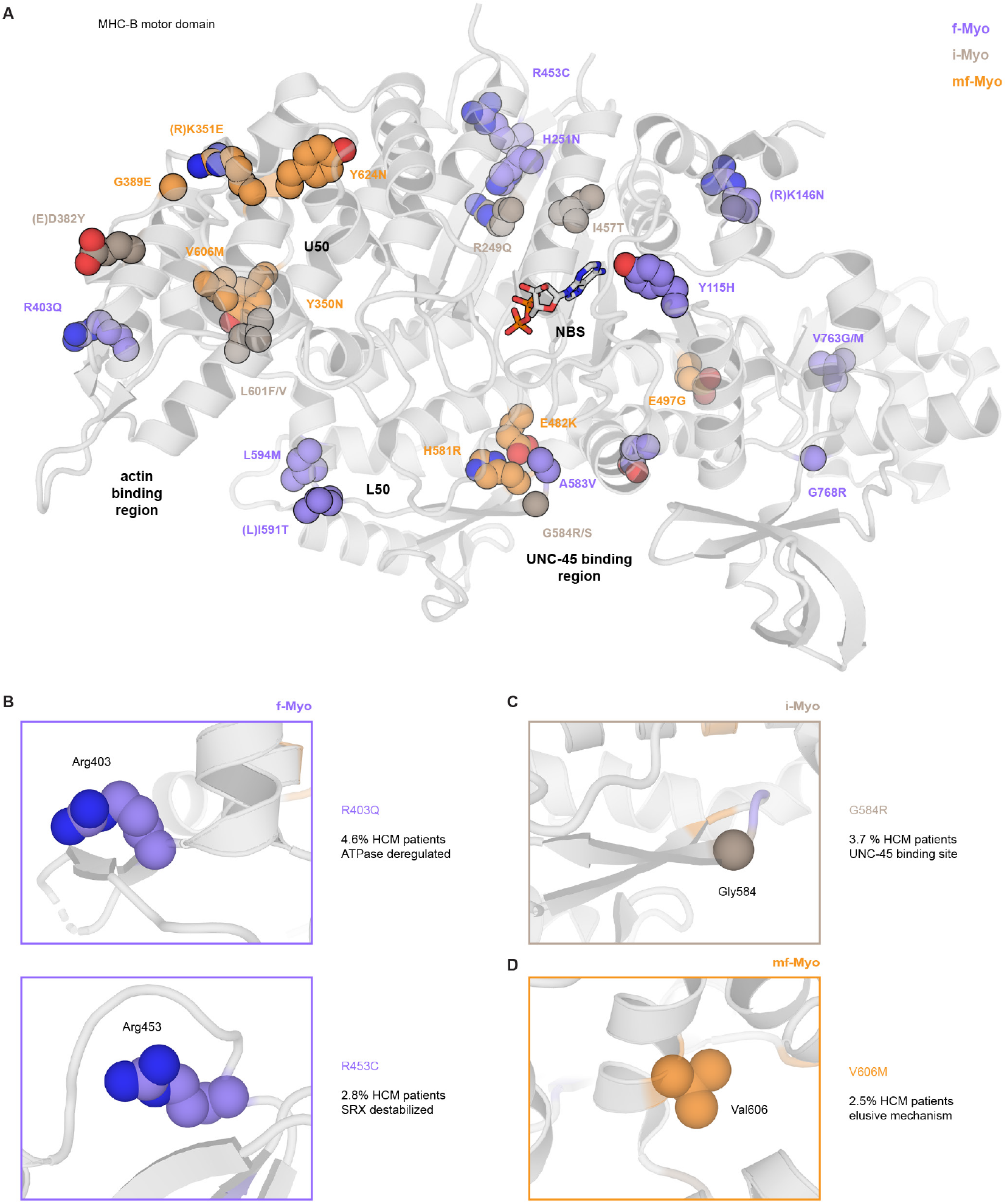
Localization of HCM mutations on HCM motor domain. **A)** Mapping of HCM mutations on MHC-B structure, color-coded according to folding defects (f-Myo, lilac; i-Myo, brown; mf-Myo, orange). **B)** Members of the f-Myo are localized on the surface of myosin (R403Q and R453C). **C)** Members of the i-Myo are localized in central parts of the motor domain implicated in ATPase activity, formation of the SRX state, or UNC-45-binding (G584R). **D)** Members of the mf-Myo are localized in residues in the inner core of the motor domain (V606M), but also in the UNC-45-binding site.

**Figure S3:**
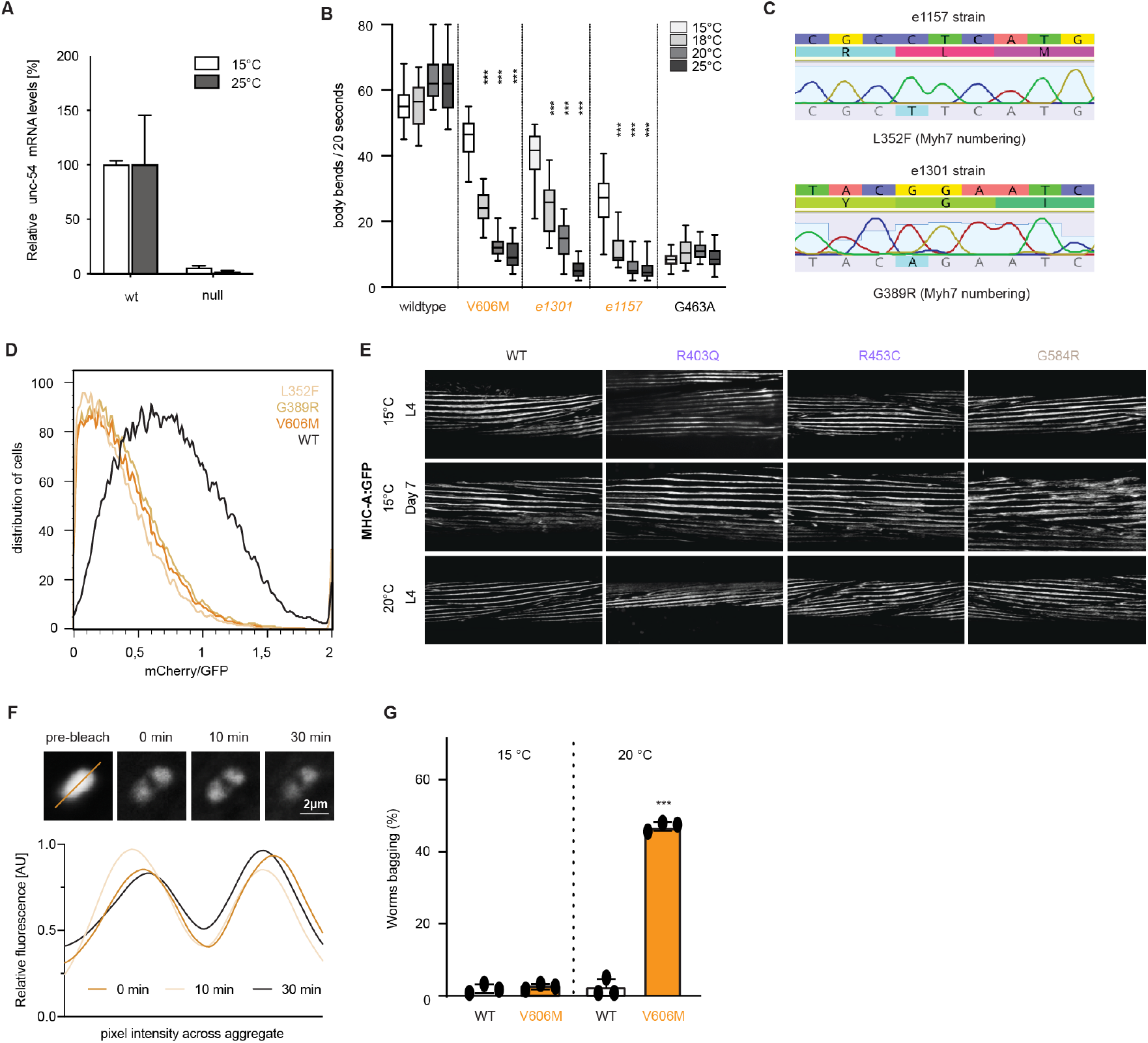
HCM mutations induce temperature-sensitive muscle defects in *C. elegans*. **A)** qPCR against *unc-54* mRNA in WT and a strain with a premature stop codon. No transcript is observed in the premature stop codon strain, probably because non-sense mediated mRNA decay. **B)** Swimming assays of ts strains and G463A inactive strain variants performed at increasing temperatures, compared to WT and V606M. **C)** Genotyping of the *C. elegans e1157* and *e1301* strains, identifying the L352F and G389R mutation, respectively. **D)** Flow-cytometry analysis of MHC-B L352F and G389R mutations, showing reduced mCherry/GFP ratio as V606M. **E)** Confocal microscopy of *C. elegans* body wall muscle using GFP labelled MHC-A myosin. Temperatures and developmental stages are indicated. **F)** Fluorescence recovery after photobleaching of V606M-GFP aggregates at 20°C. A single focal plane is shown. Relative fluorescence has been quantified by measuring relative pixel intensity across a transverse section through the aggregate and fitting a smooth curve to the data points. **G)** Expression of folding deficient V606M variant led to severe organismal dysfunctions, as represented by an increased bagging frequency.

**Figure S4:**
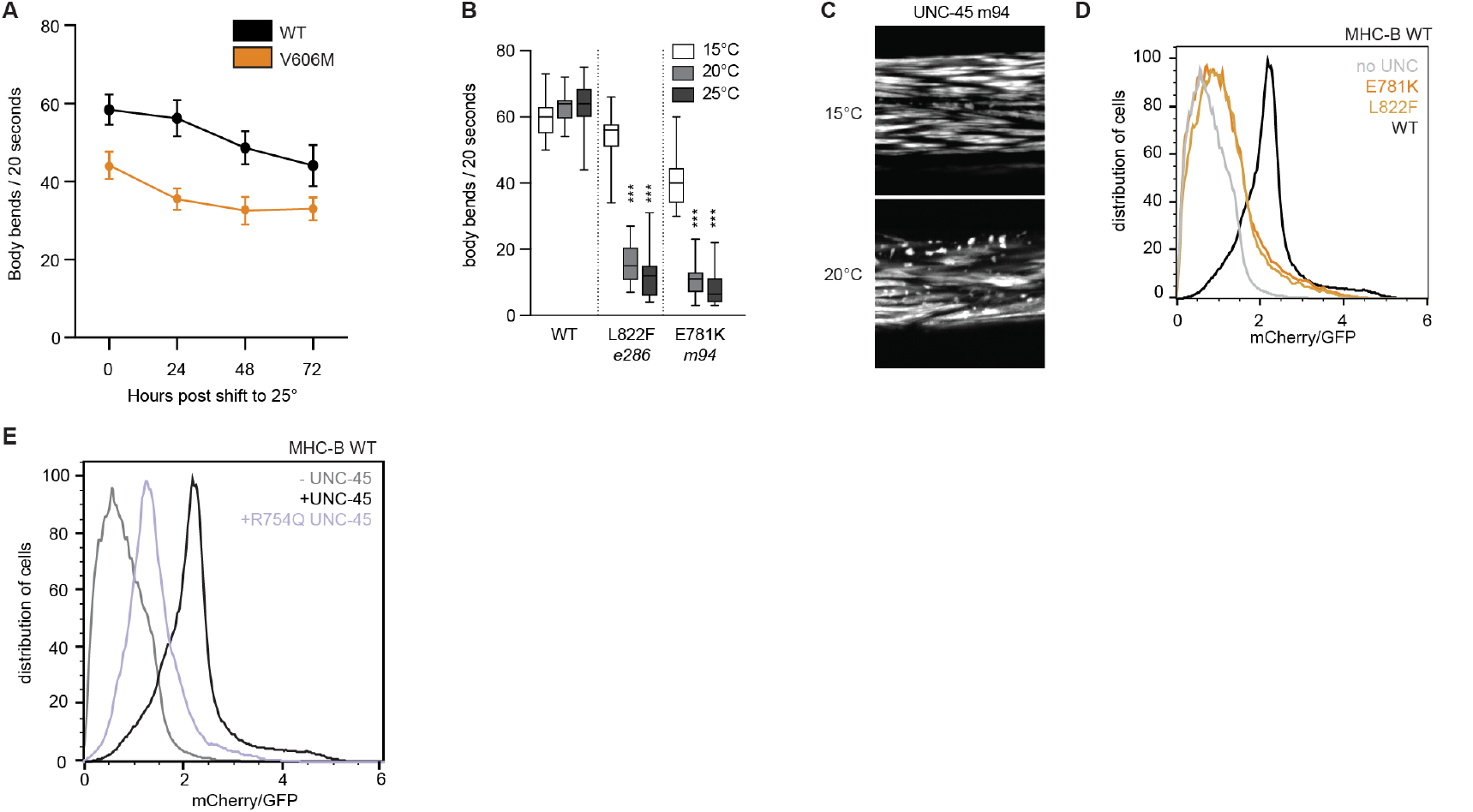
V606M destabilizes the U50 hydrophobic core of the motor domain. **A)** Swimming assays of myosin WT and V606M in adult nematodes after shifting to non-permissive temperature only after completed development. **B)** Movement of animals bearing various *unc-45(ts)* alleles, phenocopying folding-deficient MHC-B alleles regarding temperature-dependent loss of motility. **C)** Confocal microscopy of *C. elegans* (L4 stage) body wall muscle, using GFP-labelled MHC-B. The *unc-45(ts) m94* strain shows myosin aggregation. **D)** Flow-cytometry analysis of *unc-45(ts)* alleles. Co-expression with WT myosin results in low mCherry/GFP ratio in insect cells. **E)** Flow-cytometry analysis of UNC-45 R754Q with WT MHC-B, resulting in a reduction of mCherry/GFP ratio.

**Figure S5:**
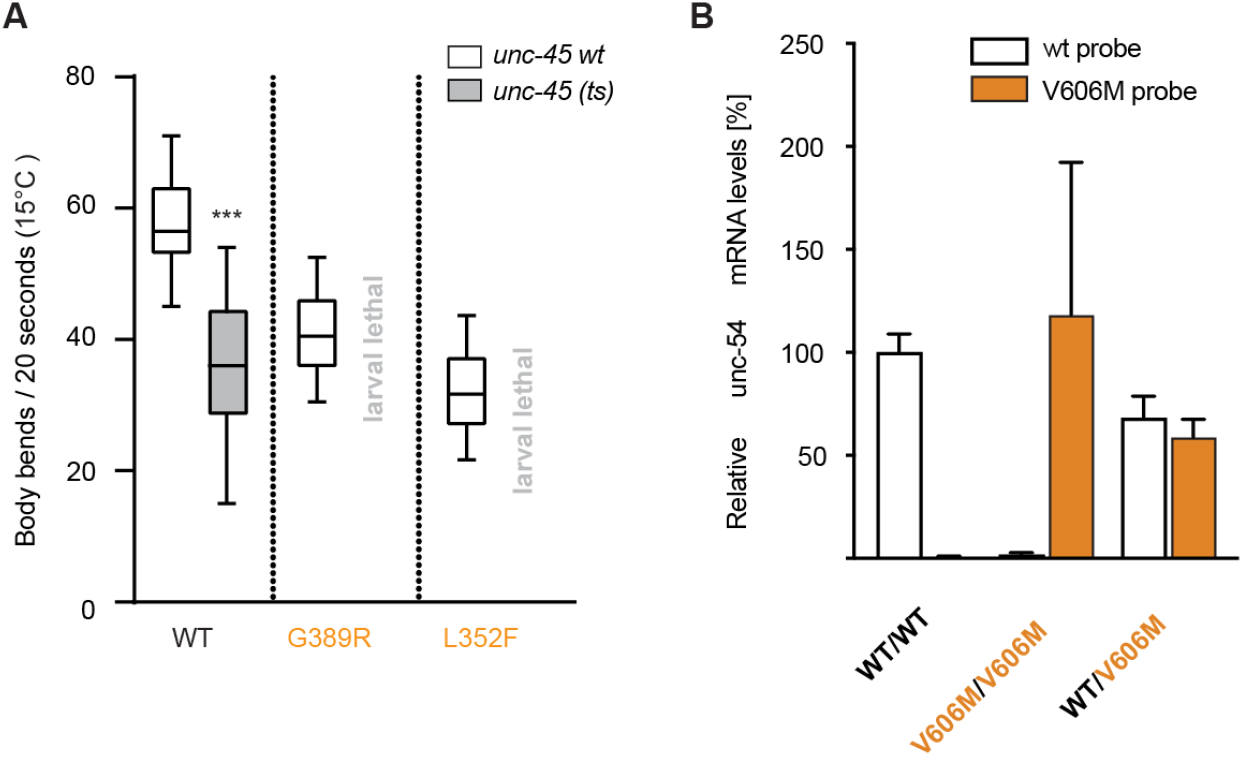
HCM mutations induce severe proteotoxic stress in muscle cells. **A)** Swimming assay monitoring synthetic effects of pathologic myosin variants and inactive UNC-45 chaperone. MHC-B ts alleles showed synthetic lethality in the *unc-45(ts)* background. **B)** Levels of *unc-54* mRNA in the heterozygous MHC-B background as measured by RT-qPCR using allele-specific TAQman probes. Heterozygotes show comparable levels of *unc-54(wt)* and *unc-54(V606M)* mRNA. n=2,3,5 respectively.

**Figure S6:**
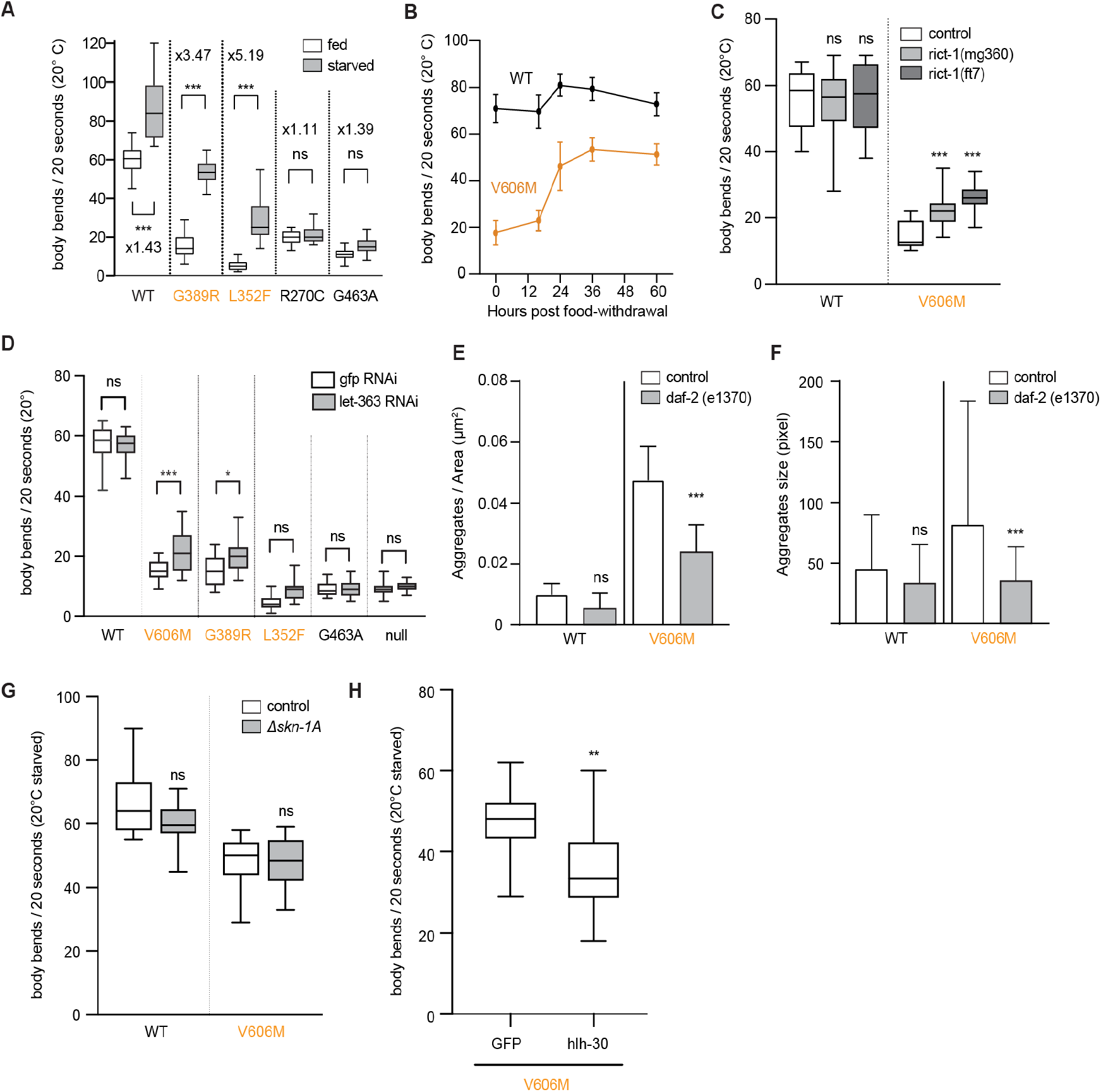
Caloric restrictions can mitigate HCM defects via autophagy. **A)** Swimming assay comparing motility of well-fed and starved worms at the L4 stage. **B)** Time course of motility after food withdrawal. The starvation rescue of folding-mutant MHC-B comes into effect after 16h and persists for a minimum of 60h. **C)** Modulation of TOR kinase activity by introducing different *rict-1* alleles resulted in a partial rescue of motility defects in V606M worms. **D)** Swimming assay of animals with RNAi against *let-363*. The folding impaired alleles are partially rescued by TOR inhibition. **E)** Quantification of aggregate number of Figure 5C. **F)** Quantification of aggregates size of Figure 5C. **G)** Swimming assay in the absence of SKN-1A in starved worms. **H)** Swimming assay silencing hlh-30 in starved worm.

**Table S1.**
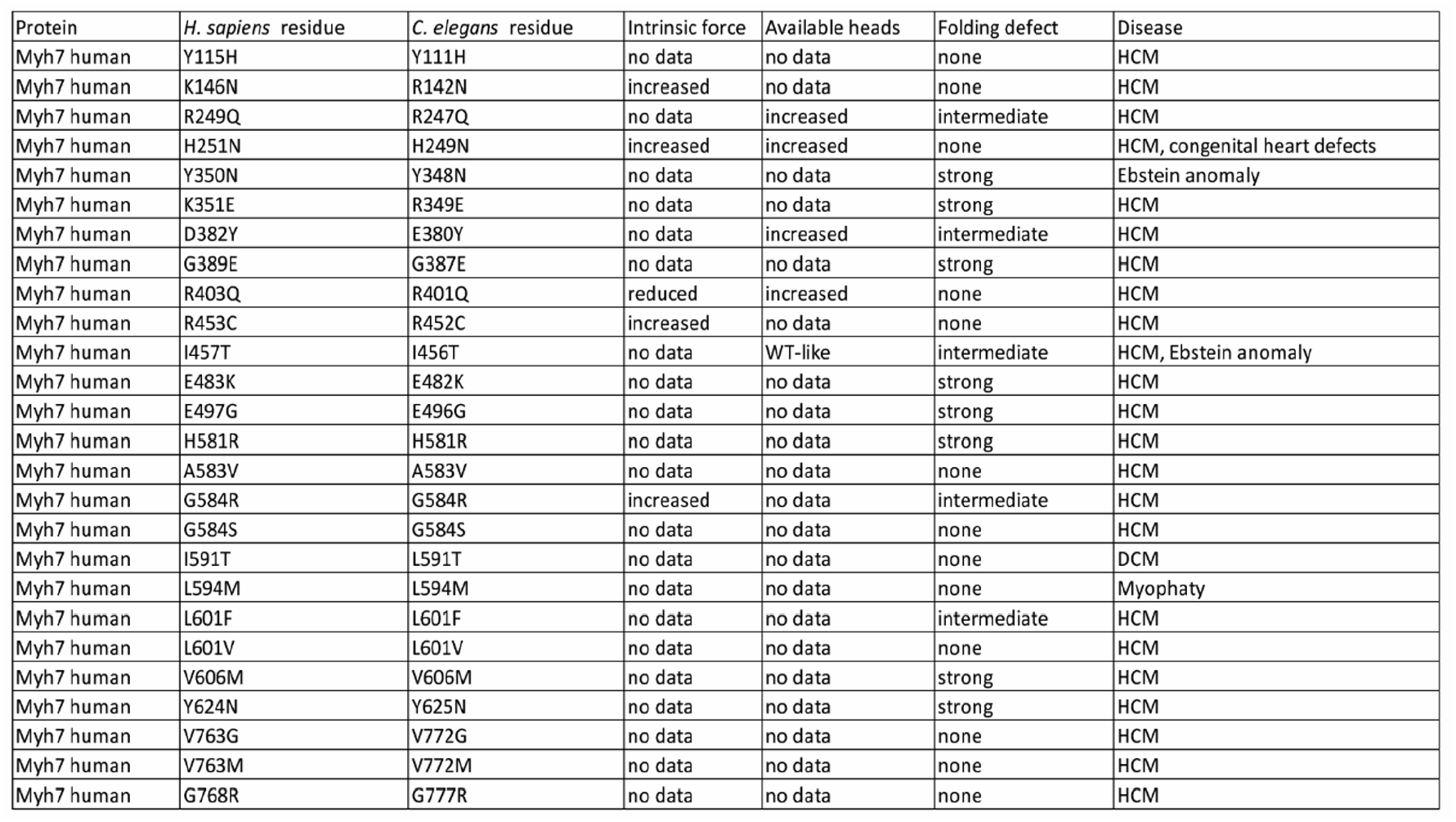
Mutations linked to HCM used in this study. Residue numbers in H. sapiens and C. elegans homologues Myh7 and MHC-B are provided. Summary of data available on intrinsic force, available heads and folding defects (this study).

