## Supplementary figures for "A fluorescent folding reporter uncovers myosin misfolding as a driver of Hypertrophic Cardiomyopathy"

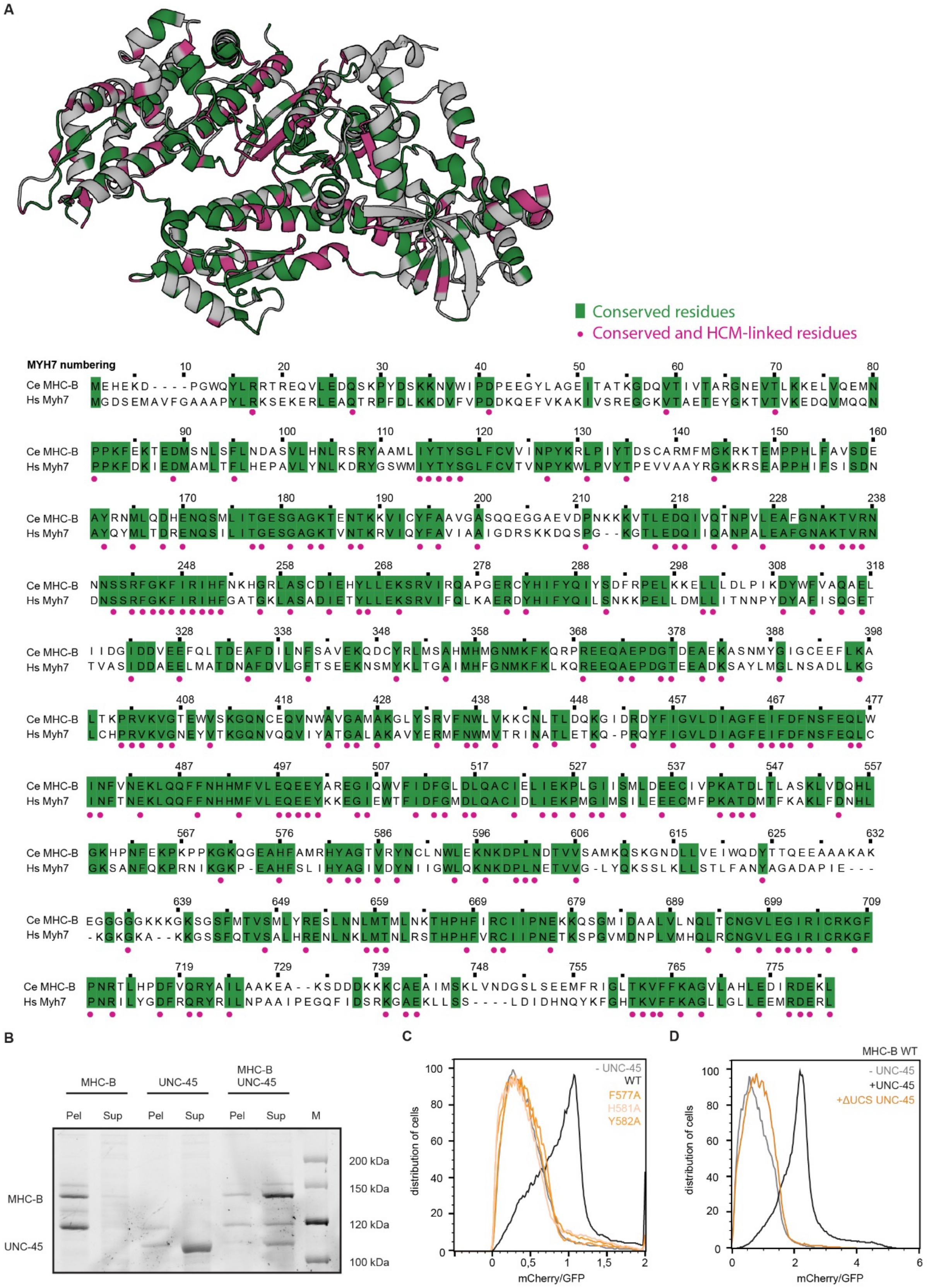

**Figure S1: HCM mutations cause graded folding defects in myosin**

**A** Alignment of *C. elegans* MHC-B to human β-cardiac myosin. Identical (conserved) residues are highlighted in green, whereas a magenta dot represents conserved residues linked to cardiomyopathies, as listed in the Human Gene Mutation Database [61]. Both conservation and disease mutations were mapped on the structure of MHC-B. **B** SDS-PAGE of soluble and insoluble fractions of MHC-B myosin folding reporter in presence or absence of UNC-45. **C** Flow-cytometry analysis of MHC mutants with WT UNC-45 **D** Flow-cytometry analysis of the ΔUCS UNC-45 mutant with WT MHC-B.

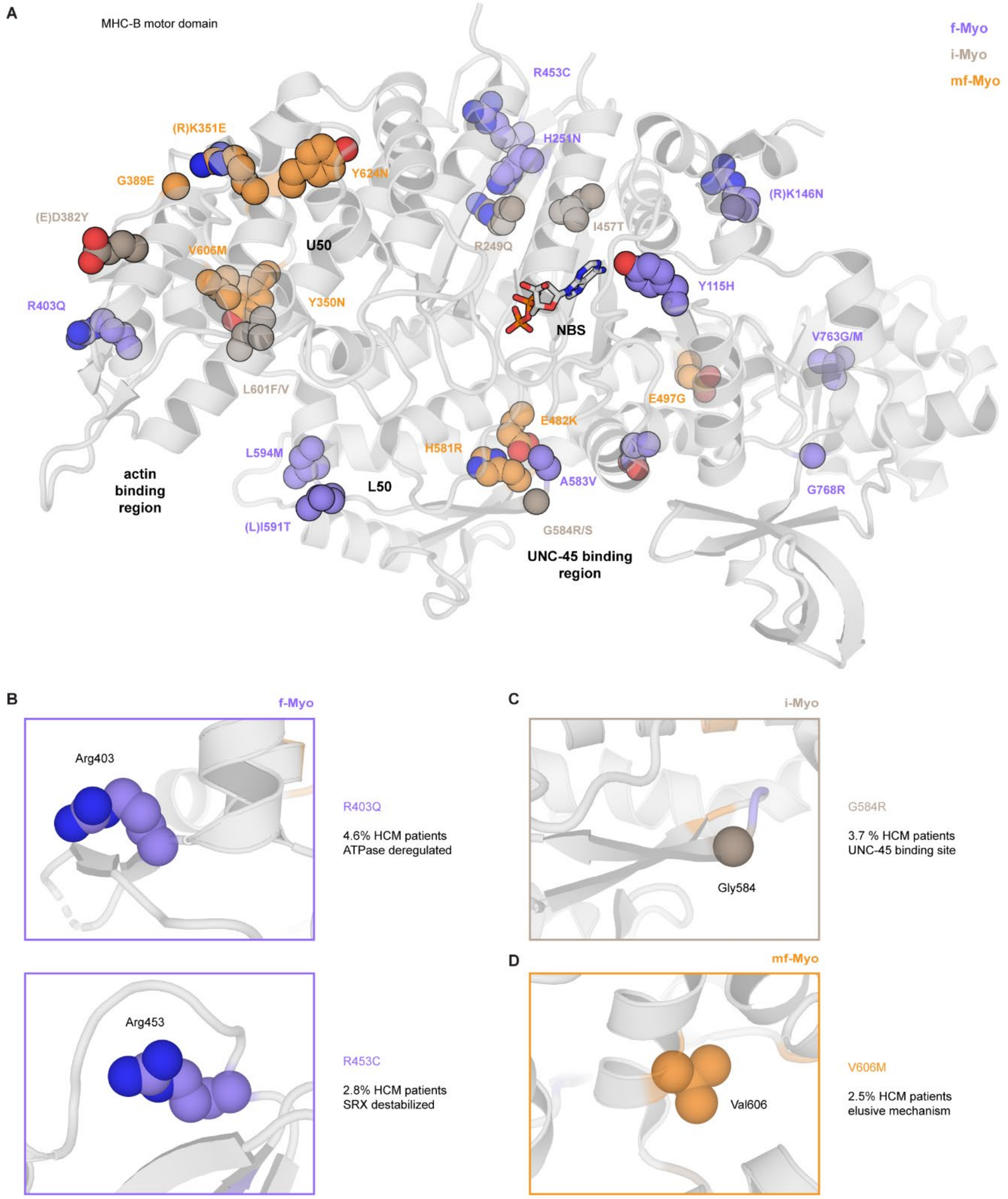


**Figure S2: Localization of HCM mutations on HCM motor domain**

**A** Mapping of HCM mutations on MHC-B structure, color-coded according to folding defects (f-Myo, lilac; i-Myo, brown; mf-Myo, orange). **B** Members of the f-Myo are localized on the surface of myosin (R403Q and R453C). **C** Members of the i-Myo are localized in central parts of the motor domain implicated in ATPase activity, formation of the SRX state, or UNC-45-

binding (G584R). **D** Members of the mf-Myo are localized in residues in the inner core of the motor domain (V606M), but also in the UNC-45-binding site.

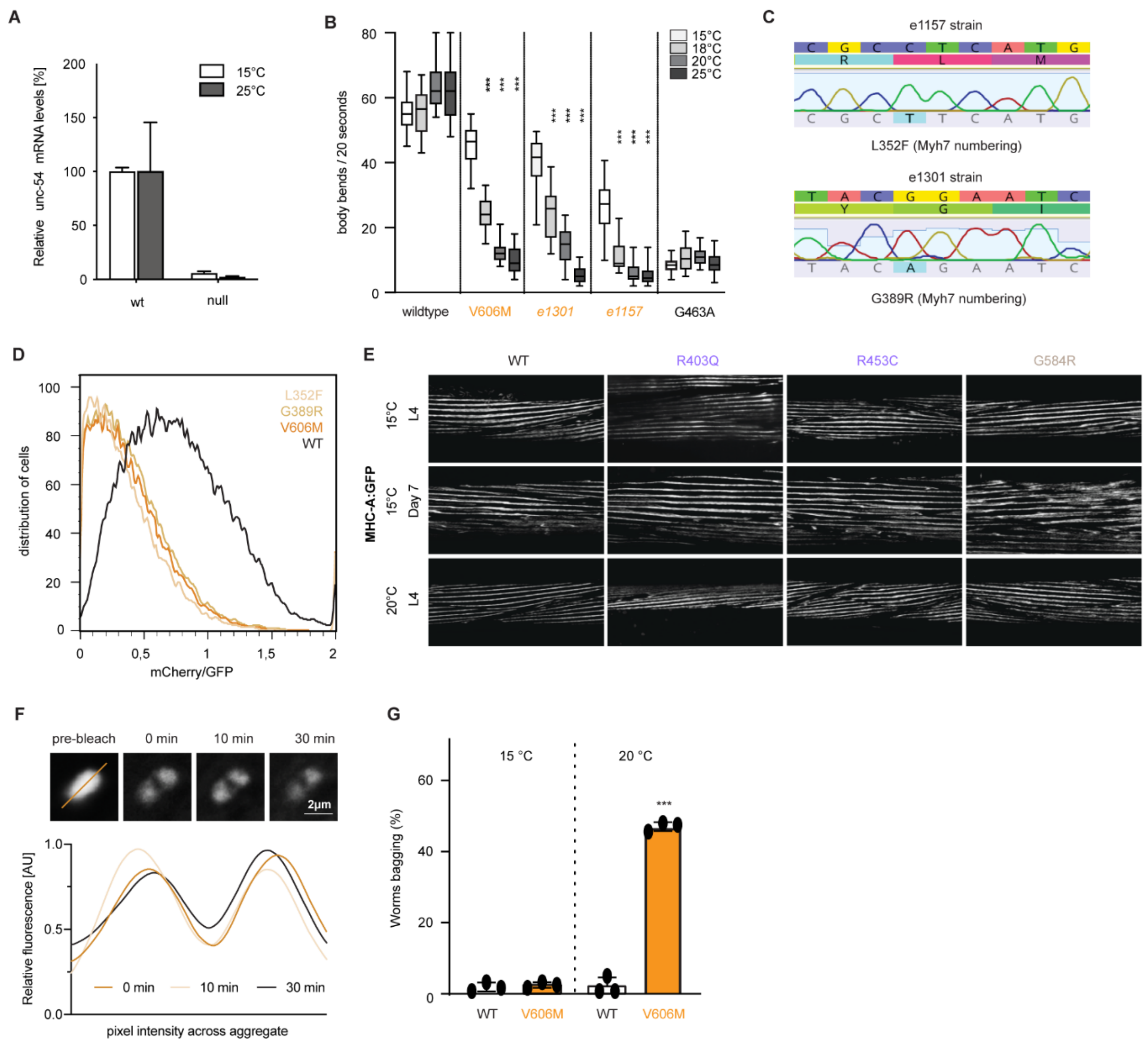


**Figure S3: HCM mutations induce temperature-sensitive muscle defects in *C. elegans***

**A** qPCR against MHC-B in WT and a strain with a premature stop codon. No transcript is observed in the premature stop codon strain, probably because non-sense mediated mRNA decay. **B** Swimming assays of ts strains and G463A inactive strain variants performed at increasing temperatures, compared to WT and V606M. **C** Genotyping of the *C. elegans e1157* and *e1301* strains, identifying the L352F and G389R mutation, respectively. **D** Flow-cytometry analysis of MHC-B L352F and G389R mutations, showing reduced mCherry/GFP ratio as V606M. **E** Confocal microscopy of *C. elegans* body wall muscle using GFP labelled MHC-A myosin. Temperatures and developmental stages are indicated. **F** Fluorescence recovery after photobleaching of V606M-GFP aggregates at 20°C. A single focal plane is shown. Relative fluorescence has been quantified by measuring relative pixel intensity across a transverse section through the aggregate and fitting a smooth curve to the data points. **G** Expression of folding deficient V606M variant led to severe organismal dysfunctions, as represented by an increased bagging frequency.

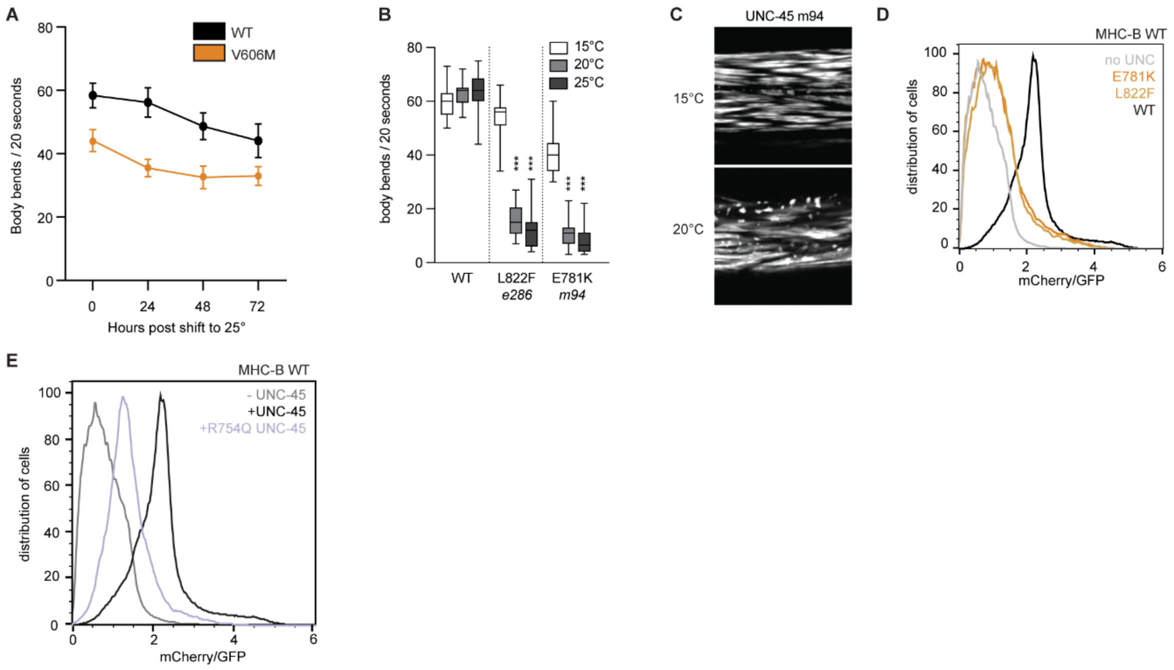


**Figure S4: V606M destabilizes the U50 hydrophobic core of the motor domain**

**A** Swimming assays of myosin WT and V606M in adult nematodes after shifting to non-permissive temperature only after completed development. **B** Movement of animals bearing various UNC-45 ts alleles, phenocopying folding-deficient MHC-B alleles regarding temperature-dependent loss of motility. **C** Confocal microscopy of *C. elegans* (L4 stage) body wall muscle, using GFP-labelled MHC-B. The UNC-45 ts strain shows myosin aggregation. **D** Flow-cytometry analysis of UNC-45 ts alleles. Co-expression with WT myosin results in low mCherry/GFP ratio in insect cells. **E** Flow-cytometry analysis of UNC-45 R754Q with WT MHC-B, resulting in a reduction of mCherry/GFP ratio.

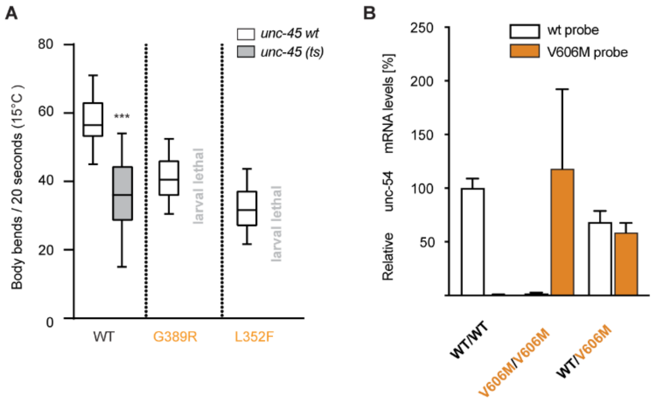


**Figure S5: HCM mutations induce severe proteotoxic stress in muscle cells**

**A** Swimming assay monitoring synthetic effects of pathologic myosin variants and inactive UNC-45 chaperone. MHC-B ts alleles showed synthetic lethality in the *unc-45(ts)* background. **B** Levels of unc-54 mRNA in the heterozygous MHC-B background as measured by RT-qPCR using allele-specific TAQman probes. Heterozygotes show comparable levels of WT and V606M mRNA. n=2,3,5 respectively.

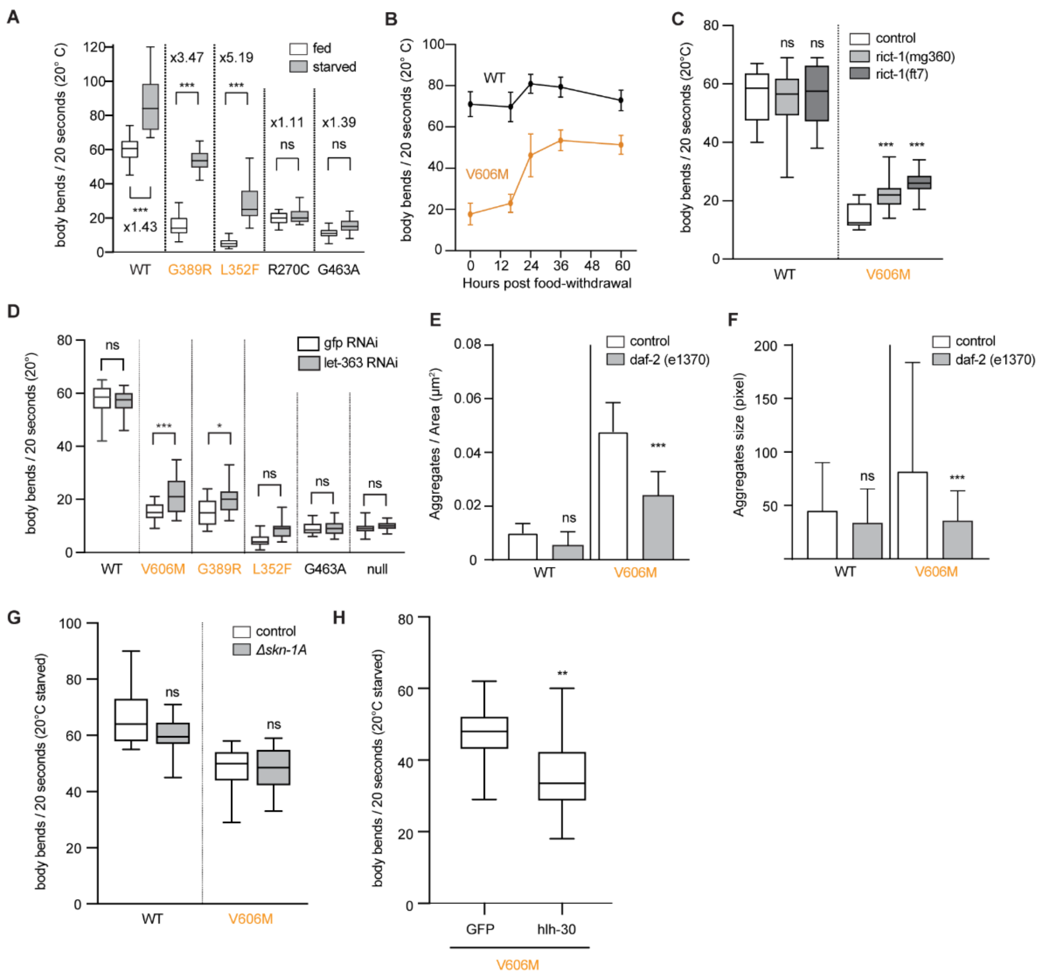


**Figure S6: Caloric restrictions can mitigate HCM defects via autophagy**

**A** Swimming assay comparing motility of well-fed and starved worms at the L4 stage. **B** Time course of motility after food withdrawal. The starvation rescue of folding-mutant MHC-B comes into effect after 16 h and persists for a minimum of 60 h. **C** Modulation of TOR kinase activity by introducing different *rict-1* alleles resulted in a partial rescue of motility defects in V606M worms. **D** Swimming assay of animals with RNAi against let363. The folding impaired alleles are partially rescued by TOR inhibition. **E** Quantification of aggregate number of Figure 5C. **F** Quantification of aggregates size of Figure 5C. **G** Swimming assay in the absence of SKN-1A in starved worms. **H** Swimming assay silencing hlh-30 in starved worm.

| Protein | *H. sapiens* residue | *C. elegans* residue | Intrinsic force | Available heads | Folding defect | Disease |
|---|---|---|---|---|---|---|
| Myh7 human | Y115H | Y111H | no data | no data | none | HCM |
| Myh7 human | K146N | R142N | increased | no data | none | HCM |
| Myh7 human | R249Q | R247Q | no data | increased | intermediate | HCM |
| Myh7 human | H251N | H249N | increased | increased | none | HCM, congenital heart defects |
| Myh7 human | Y350N | Y348N | no data | no data | strong | Ebstein anomaly |
| Myh7 human | K351E | R349E | no data | no data | strong | HCM |
| Myh7 human | D382Y | E380Y | no data | increased | intermediate | HCM |
| Myh7 human | G389E | G387E | no data | no data | strong | HCM |
| Myh7 human | R403Q | R401Q | reduced | increased | none | HCM |
| Myh7 human | R453C | R452C | increased | no data | none | HCM |
| Myh7 human | I457T | I456T | no data | WT-like | intermediate | HCM, Ebstein anomaly |
| Myh7 human | E483K | E482K | no data | no data | strong | HCM |
| Myh7 human | E497G | E496G | no data | no data | strong | HCM |
| Myh7 human | H581R | H581R | no data | no data | strong | HCM |
| Myh7 human | A583V | A583V | no data | no data | none | HCM |
| Myh7 human | G584R | G584R | increased | no data | intermediate | HCM |
| Myh7 human | G584S | G584S | no data | no data | none | HCM |
| Myh7 human | I591T | L591T | no data | no data | none | DCM |
| Myh7 human | L594M | L594M | no data | no data | none | Myophaty |
| Myh7 human | L601F | L601F | no data | no data | intermediate | HCM |
| Myh7 human | L601V | L601V | no data | no data | none | HCM |
| Myh7 human | V606M | V606M | no data | no data | strong | HCM |
| Myh7 human | Y624N | Y625N | no data | no data | strong | HCM |
| Myh7 human | V763G | V772G | no data | no data | none | HCM |
| Myh7 human | V763M | V772M | no data | no data | none | HCM |
| Myh7 human | G768R | G777R | no data | no data | none | HCM |

**Table S1**

Mutations linked to HCM used in this study. Residue numbers in *H. sapiens* and *C. elegans* homologues Myh7 and MHC-B are provided. Summary of data available on intrinsic force, available heads and folding defects (this study).
